# The Genetic Basis of phage susceptibility, cross-resistance and host-range in *Salmonella*

**DOI:** 10.1101/2020.04.27.058388

**Authors:** Benjamin A. Adler, Alexey E. Kazakov, Crystal Zhong, Hualan Liu, Elizabeth Kutter, Lauren M. Lui, Torben N. Nielsen, Heloise Carion, Adam M. Deutschbauer, Vivek K. Mutalik, Adam P. Arkin

## Abstract

Though bacteriophages (phages) are known to play a crucial role in bacterial fitness and virulence, our knowledge about the genetic basis of their interaction, cross-resistance and host-range is sparse. Here, we employed genome-wide screens in *Salmonella enterica* serovar Typhimurium to discover host determinants involved in resistance to eleven diverse lytic phages including 4 new phages isolated from a therapeutic phage cocktail. We uncovered 301 diverse host factors essential in phage infection, many of which are shared between multiple phages demonstrating potential cross-resistance mechanisms. We validate many of these novel findings and uncover the intricate interplay between RpoS, the virulence-associated general stress response sigma factor and RpoN, the nitrogen starvation sigma factor in phage cross-resistance. Finally, the infectivity pattern of eleven phages across a panel of 23 genome sequenced *Salmonella* strains indicates that additional constraints and interactions beyond the host factors uncovered here define the phage host range.

## Introduction

There is increasing evidence that bacteriophages are a critical feature of microbial ecology, evolution, virulence and fitness(Abedon, 2009; Breitbart and Rohwer, 2005; Koskella and Taylor, 2018; Shkoporov and Hill, 2019; Suttle, 2007). However, knowledge about the molecular and genetic determinants of host-phage interactions and how they vary across the populations of both are sparse even in otherwise well-studied model systems(Brüssow, 2013; de Jonge et al., 2019; Nobrega et al., 2018; Rostøl and Marraffini, 2019; Samson et al., 2013; Weitz et al., 2013; Young and Gill, 2015). This derives, in part, from the technological limitations to resolve the incredible specificity and the complex suite of bacterial mechanisms that confer both resistance and sensitivity to the phage(De Smet et al., 2017; Nobrega et al., 2018; Young and Gill, 2015). A given bacterial host is likely to be susceptible to multiple phages, while a given phage may infect a specific array of hosts and their variants. There are few, if any, studies that map these mechanisms across phages and hosts, their interdependencies, and how variations in these mechanisms encode tradeoffs in host and phage fitness under different conditions(Calendar, 2012; Casjens and Hendrix, 2015; De Smet et al., 2017; Karam and Drake, 1994; Molineux, 2002). However, this information is critical to an understanding of microbial ecology and possibly exploiting the predator-prey dynamics for applications(Campbell, 2003; Díaz-Muñoz and Koskella, 2014; Lenski, 1988; Mirzaei and Maurice, 2017).

Knowledge of phage susceptibility and resistance determinants underlies a number practical applications of phages. These applications span the use of phages and their combinations as biocontrol agents to improve water quality, decontaminate food, protect agricultural yield, and defend and improve human health(Gordillo Altamirano and Barr, 2019; Keen and Adhya, 2015; Kortright et al., 2019; Pirnay and Kutter, 2020). For example, because of the apparent ubiquity of lytic phage with high host specificity for nearly any known pathogenic bacterial strain, phages may provide a powerful alternative or adjutant to antibiotic therapies. The development of such therapeutic phage formulations is pressing due to the alarming rise of antibiotic resistance(Gordillo Altamirano and Barr, 2019; Keen and Adhya, 2015; Kortright et al., 2019; Young and Gill, 2015). By characterizing the genetic basis of a bacterium’s susceptibility and resistance to a given phage and the pattern of cross-resistance or cross-sensitivity with other phages, we can uncover evolutionary trade-offs on bacteria-phage interactions. These insights could also identify knowledge-gaps in our understanding of the host-range of a phage and offer therapeutic solutions to recalcitrant infections(Chan et al., 2016; Gordillo Altamirano et al., 2021; Mangalea and Duerkop, 2020; Shin et al., 2012; Trudelle et al., 2019; Wright et al., 2018). For instance, by leveraging phages that target different receptors, combinations of phages or phage cocktails can be rationally formulated to both extend the host range and limit the rate of resistance emergence(Bai et al., 2019; Chan et al., 2013; Kortright et al., 2019; Tanji et al., 2004; Yen et al., 2017). Such strategies can be further augmented by selecting phages that specifically bind to bacterial virulence or antibiotic resistance factors to benefit from evolutionary trade-offs in rational therapeutic outcomes (Chan et al., 2013; Kortright et al., 2019).

Compared to other antimicrobials, characterization of infectivity and cross-resistance between a panel of phages has been limited to a few model organisms and remained phenomenological until recently(Hudson et al., 1978; Lindberg and Hellerqvist, 1971; Marti et al., 2013; Samuel et al., 1999; Tu et al., 2017; Wright et al., 1980). The advent of genome-wide saturated transposon sequencing (Bohm et al., 2018; Chan and Turner, 2020; Christen et al., 2016; Cowley et al., 2018; Pickard et al., 2013) and the corresponding DNA bar-code based modifications has enabled the high-throughput and low cost genome-scale screening for the genetic determinants of these phenomena(Carim et al., 2020; Mutalik et al., 2019, 2020; Rousset et al., 2018; Wetmore et al., 2015). Since this economically permits the independent screening of many phages against a host library, it is now possible to determine and compare the genes that affect the successful infection of one or more phages. These insights further suggest possible mechanisms of cross-resistance (ie single mutations that confer resistance to multiple phages) and collateral-sensitivity (ie single mutations that cause resistance to one phage while sensitizing to another phage) that might arise when the host is naturally exposed to different combinations of these phages in the environment. As an example of this approach, we recently employed a high-throughput genetic screening platform to characterize the phage resistance landscape in *Escherichia coli* (*E*.*coli*) at an unprecedented scale(Mutalik et al., 2020). However, the scale and benefits of these technologies have not yet been realized outside of such model organisms, where bacterial physiology and phage-host interactions can be dramatically different. Here, we employ a genome-wide loss-of-function screening technology to discover the genetic determinants of phage susceptibility in *Salmonella enterica, a* globally important infectious bacteria whose variants are responsible for the vast majority of bacterial food-borne infections with an annual cost of $3.7B dollars in 2013(Maculloch et al., 2015). Though *Salmonella enterica* serovar Typhimurium (*S*. Typhimurium) has been used in the past as a model host to study phage infections(Graña et al., 1985; Lee et al., 2013; Lindberg and Hellerqvist, 1971; MacPhee et al., 1975; Marti et al., 2013; Schwartz, 1980; Wilkinson et al., 1972; Wright et al., 1980), most *Salmonella* phages including therapeutically employed phage formulations have limited characterization regarding target receptors and host resistance mechanisms (Bai et al., 2019; Gao et al., 2020; Islam et al., 2019; McCallin et al., 2018; Petsong et al., 2019; Zschach et al., 2015). With increased numbers of *Salmonella* infections that are resistant to antibiotics and displaying increased virulence(Medalla et al., 2016; (u.s.) and Centers for Disease Control and Prevention (U.S.), 2019), it is imperative to characterize phages and their resistance patterns to enable their rational use as diagnostic and antimicrobial agents.

In this study, we use a *S. enterica* serovar Typhimurium (*S*. Typhimurium) LT2 derivative that serves as a genetically-accessible model for a clade of *Salmonella* responsible for food-borne enteric infections in humans. Using a barcoded transposon mutant library, we identified bacterial genes whose loss confers altered sensitivity to 11 diverse double-stranded DNA phages including 4 new phages isolated from a therapeutic phage cocktail. Our screens identified known (and proposed new) receptors and also yielded novel host factors important in phage infection, some of which we validate using single-gene deletion strains. Our genetic analysis allowed high resolution mapping of phage interactions with diverse cell surface components and the operation of specific global regulatory systems that mediate specific metabolisms (e.g. *rpoN*) or virulence and stress response (e.g. *rpoS*). The diversity of cellular processes that influence sensitivity to phage suggest that there may be multiple routes for cross-resistance and cross-sensitivity to emerge due to phage exposure in the environment or when used as therapeutic cocktails. Finally, to assess if phage susceptibility and phage host-range can be predicted based on the genetic determinants uncovered by our screens, we measured and analyzed phage infectivity against a panel of 23 *S*.Typhimurium strains representative of naturally occurring genetic diversity and phage infectivity.

## Results

### Identifying *Salmonella* genes involved in resistance to 11 diverse phages

We previously established a high-throughput approach to assay gene fitness using genome-wide, random barcoded transposon sequencing (RB-TnSeq) (Price et al., 2018; Wetmore et al., 2015). To systematically characterize phage infectivity pathways in *Salmonella*, we first constructed an RB-TnSeq library in *S*. Typhimurium LT2 derivative strain MS1868 (Graña et al., 1985) (Methods). As an LT2-derived strain, *S*. Typhimurium MS1868 benefits from a long history of *Salmonella* phage-host genetic interaction studies and well-characterized phage-resistant genotypes (Hudson et al., 1978; Lindberg and Hellerqvist, 1971; Marti et al., 2013; Samuel et al., 1999; Tu et al., 2017; Wilkinson et al., 1972; Wright et al., 1980). In addition, *S*. Typhimurium MS1868 is a restriction-minus genetic background (Graña et al., 1985), which would potentially help uncover additional phage resistance factors by expanding the number of phages infectious to the RB-TnSeq library. After transposon mutagenesis, we obtained a 66,996 member pooled library consisting of transposon-mediated disruptions across 3,759 of 4,610 genes, with an average of 14.8 disruptions per gene (median 12) (Figure 1A). We note that our library does not have sufficient coverage of some likely non-essential genes that are likely to play an important role in phage infection, for example igaA(Cho et al., 2014; Mariscotti and García-del Portillo, 2009; Mutalik et al., 2020). Additional details for the composition of the *S*. Typhimurium MS1868 library and comparison to a related single-gene-deletion library(Porwollik et al., 2014) can be found in Table S1 and Dataset S1.

**Figure 1.**
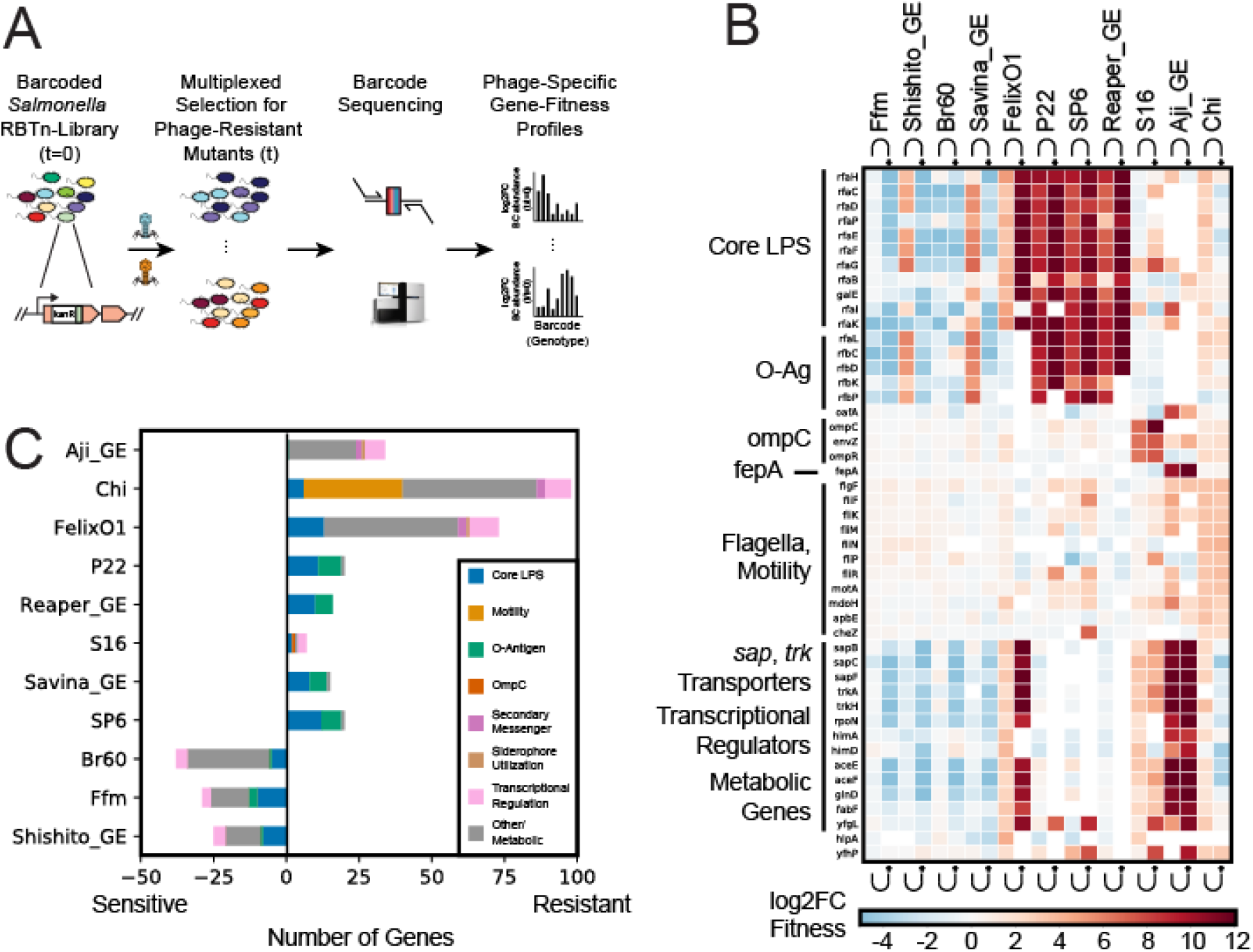
Genome-wide screen to identify host factors involved in phage infection. (A) Overview of pooled fitness assays. For additional details, see methods. Briefly, for each experiment, *S*. Typhimurium RB-TnSeq library was exposed to a high MOI of one of eleven dsDNA *Salmonella* phages. Strains were tracked by quantifying the abundance of DNA barcodes associated with each strain by Illumina sequencing. Phage-specific gene fitness profiles were calculated by taking the log2-fold-change of barcode abundances post-(t) to pre-(t=0) phage predation. High fitness scores indicate that loss of genetic function in *Salmonella* confers fitness against phage predation. (B) Heatmap of top 10 high-confidence gene scores per phage are shown (many genes are high-confidence hits to multiple phages). Both planktonic and solid plate data are shown. Three rough-LPS binding phages Br60, Ffm, and Shishito_GE do not infect wild-type MS1868, but can infect specific MS1868 mutants, overall showing negative fitness fitness values in our screen. Noncompetitive, solid agar growth experiments are marked with a (*). (C) Total number of high-scoring genes per phage and their functional role. Input data for Figures 1B and 1C is found in Dataset S4 and can be recreated using Supplementary Code - Figure1BC.

We collected 11 lytic, dsDNA *Salmonella* phages, some of which are currently employed in therapy and diagnostics. These phages are diverse, representing 5 of the 9 major dsDNA phage families currently listed by the International Committee on Taxonomy of Viruses (ICTV): 3 from Myoviridae (FelixO1, S16, and Savina_GE), 1 from Podoviridae (P22), 4 from Autographiviridae (Br60, Ffm, Shishito_GE, and SP6), 2 from Siphoviridae (Chi, and Reaper_GE), and 1 from Demerecviridae (Aji_GE). Though the receptors for 4 of the phages investigated here, Chi, FelixO1, P22 (obligately lytic mutant), and S16 are relatively well-studied, (Graña et al., 1985; Lee et al., 2013; Lindberg and Hellerqvist, 1971; MacPhee et al., 1975; Marti et al., 2013; Schwartz, 1980; Wilkinson et al., 1972; Wright et al., 1980), only P22 phage has been subjected to a genome-wide genetic screen (Bohm et al., 2018). Additionally, Br60, Ffm, and SP6 have suspected host-factor requirements for their infectivity cycle but otherwise have not been extensively studied (Gebhart et al., 2017; Lindberg and Hellerqvist, 1971; Tu et al., 2017). This panel of 11 phages also consists of 4 newly isolated phages from a commercial phage-cocktail preparation from the Republic of Georgia (see Methods, Dataset S7): Aji_GE_EIP16 (Aji_GE), Reaper_GE_8C2 (Reaper_GE), Savina_GE_6H2 (Savina_GE), and Shishito_GE_6F2 (Shishito_GE). All 11 phages except 3 (Br60, Ffm and Shishito_GE) infect wild-type *S*. Typhimurium LT2 MS1868, which has an intact O-antigen (known as ‘smooth-LPS’). Br60, Ffm and Shishito_GE phages were grown using a ‘rough-LPS’ mutant strain of *Salmonella* which consists of only core LPS (see Methods). Thus, the phage panel used here contains phages that either bind to smooth or rough LPS strains and allows comparison of the key host factors important in their infectivity cycles.

To identify *Salmonella* genes important for phage infection, we challenged the *S*. Typhimurium mutant library with each of the 11 dsDNA lytic phages (Table 1) at multiplicities of infection ≥2 in both planktonic and non-competitive solid plate fitness experiments, and collected the surviving phage-resistant strains post incubation (Figure 1A, Methods). From samples collected before and after phage incubation, we sequenced the 20 base pair DNA barcodes (i.e. BarSeq) associated with each transposon mutant. We then calculated strain and gene fitness scores as the relative log2-fold-change of barcode abundances before versus after phage selection, as previously described (Mutalik et al., 2020; Price et al., 2018; Wetmore et al., 2015) (Figure 1A, Methods). Thus, in this study, a high positive fitness score indicates loss-of-function mutants in *Salmonella* that are resistant to phage infection. We observed very strong phage selection pressures during these competitive fitness experiments, consistent with our earlier observations (Mutalik et al., 2020), and thus we mostly limited our analysis to positive fitness scores. As expected with our MS1868 library primarily consisting of O-antigen positive mutants, the vast majority of gene disruptions in MS1868 showed no significant fitness benefit against rough-LPS binding Br60, Ffm and Shishito_GE phages. However, we noticed strong fitness defects in many of the LPS and O-antigen mutants in our library (Figure 1BC, Dataset S4), consistent with optimal adsorption and infection in O-antigen-defective *Salmonellae*.

**Table 1.** Bacteriophages employed in this study. Virus families were assigned via ICTV taxonomy release 2019. For new phages isolated in this study, the family of the nearest BLASTN relative was reported (in line with ICTV 2019 standards). This information can be found in Dataset S7.

| Phage | Family | Established Receptor? | Source | Reference |
| --- | --- | --- | --- | --- |
| Aji_GE_EIP16 (Aji_GE) | Demerecviridae | No | Intesti Bacteriophage formulation M2-601 | This study. |
| Br60 | Autographiviridae | "Rough LPS *Salmonella*" | *Salmonella* Genetic Stock Center (SGSC) | (Wilkinson et al., 1972) |
| Chi | Siphoviridae | Flagella | Gift from Jason Gill | (Samuel et al., 1999; Schade et al., 1967) |
| FelixO1 | Myoviridae | Outer core LPS | Félix d'Hérelle Reference Center for Bacterial Viruses | (Gebhart et al., 2017; Lindberg and Hellerqvist, 1971; Tu et al., 2017) |
| Ffm | Autographiviridae | "Rough-LPS *Salmonella*" | *Salmonella* Genetic Stock Center (SGSC) | (Graña et al., 1985; Lee et al., 2013; Lindberg and Hellerqvist, 1971; MacPhee et al., 1975; Marti et al., 2013; Schwartz, 1980; Wilkinson et al., 1972; Wright et al., 1980) |
| P22 | Podoviridae | O-Antigen LPS | Gift from Richard Calendar | (Bohm et al., 2018; Steinbacher et al., 1997; Wright et al., 1980) |
| Reaper_GE_8C2 (Reaper_GE) | Siphoviridae | No | Intesti Bacteriophage formulation M2-601 | This study |
| S16 | Myoviridae | OmpC | Félix d'Hérelle Reference Center for Bacterial Viruses | (Marti et al., 2013) |
| Savina_GE_6H2 (Savina_GE) | Myoviridae | No | Intesti Bacteriophage formulation M2-601 | This study |
| Shishito_GE_6F2 (Shishito_GE) | Autographiviridae | No | Intesti Bacteriophage formulation M2-601 | This study |
| SP6 | Autographiviridae | O-Antigen LPS | Gift from Ian J Molineaux | (Tu et al., 2017; Wright et al., 1980) |

In aggregate, we performed 42 genome-wide RB-TnSeq assays across liquid and solid growth formats and discovered 301 phage-gene interactions (with 184 unique gene hits) that are important for phage infection across the 11 phages studied (Figure 1BC, Datasets S2 and S4). Though solid plate assay results were largely consistent with planktonic assays, some resulted in genes with stronger fitness effects. Across all fitness experiments, we observed at least one gene with a high fitness score (except 3 phages that infect rough-LPS strains), affirming the successful competitive growth of mutants under phage selection. Some *Salmonella* phages show enrichment of strains with disruptions in multiple genes, while other phages enrich strains with disruptions in a more limited number of genes (Figure 1B). For example, we observed 98 genes enriched after Chi phage challenge and 73 high-scoring genes after Felix O1 challenge, yet only 7 high-scoring genes after S16 phage challenge. As expected in any phage selection experiment, we observed enrichment of genes that encode components of the cell envelope. Nonetheless, we also identified dozens of genes that encode cytoplasmic components not previously associated with phage resistance. To further categorize the genetic basis of phage resistance, we manually classified all identified genes with high fitness values into broad-functional categories: core-LPS and O-antigen biosynthesis, motility, secondary messengers, transcription factors and other metabolism (Figure 1C). These results demonstrate that genes downstream from phage receptors are important for phage infectivity.

### Both receptor and non-receptor host factors are involved in phage infection

A key-determining step in the phage infectivity cycle is the interaction of phages with any bacterial cell surface-exposed molecules or receptors. Consequently, any changes in the structure or level of these surface-exposed molecules that accompany resistance to specific phages are usually assigned a function of phage receptor. To confirm the effectiveness of our genetic screen, we looked for receptors that are known for a few of the phages used in this work (Bohm et al., 2018; Hudson et al., 1978; Lindberg and Hellerqvist, 1971; MacPhee et al., 1975; Marti et al., 2013; Samuel et al., 1999). Indeed, in agreement with published data available for FelixO1, P22, Chi, SP6 and S16 phages, we found high fitness scores for candidate receptor genes with >1,000 fold enrichment of transposon mutants. These included genes encoding protein receptors such as *ompC* (outer membrane porin C) for S16 and flagellar body for Chi phage, while LPS and O-antigen biosynthesis genes for P22, SP6 and FelixO1 phages (Figure 1C). Our results are also largely consistent with a recent genome-wide screen in *Salmonella* against P22 infection (Bohm et al., 2018). Though O-antigen and outer core GlcNAc (the biosynthetic product of RfaK) have been known as SP6 and as FelixO1 phage receptors respectively (Hudson et al., 1978; Lindberg and Hellerqvist, 1971; Marti et al., 2013; Samuel et al., 1999; Tu et al., 2017; Wilkinson et al., 1972; Wright et al., 1980), our genome-wide screens provided an array of additional, non-receptor genes as target loci for phage resistance selection. A detailed description and analysis of outer membrane components such as LPS required for these phages can be found in Text S1.

In addition to the genes coding for phage receptors, our genetic screens also uncovered high-scoring genes that are known to be involved in the regulation of target receptors. For example, deletion of the EnvZ/OmpR two component system involved in the regulation of *ompC* and gene products involved in the regulation of cellular motility (*nusA, tolA, cyaA* and guanosine penta/tetraphosphate ((p)ppGpp) biosynthesis and metabolism all showed high fitness scores in the presence of S16 and Chi phages, respectively(Graña et al., 1985; Lee et al., 2013; Lindberg and Hellerqvist, 1971; MacPhee et al., 1975; Marti et al., 2013; Schwartz, 1980; Wilkinson et al., 1972; Wright et al., 1980). These high-scoring gene candidates were previously not known to be associated with phage resistance in *Salmonella*. Other than the phages mentioned above that bind to surface components of smooth-LPS *Salmonellae*, we also screened Br60 and Ffm phages, which are known to strictly infect rough-LPS strains and not bind to smooth WT MS1868 parental strain (as O-antigen structure probably occludes their native receptor)(Gebhart et al., 2017; Lindberg and Hellerqvist, 1971; Tu et al., 2017). As our MS1868 library primarily consists of O-antigen positive mutants, the vast majority of gene disruptions in MS1868 showed no significant fitness benefit against these phages. However, we noticed strong negative scores for many of the LPS and O-antigen mutants in our library (Dataset S4), indicating these strains have rough LPS phenotype and are sensitive to Br60 and Ffm phages. As an additional resource, we determined the specific rough-LPS requirements for these phages using an O-antigen deficient library and individual mutant susceptibility assays (Text S1 - Extended Results: Uncovering Host-Factors of Rough-LPS Requiring Phages Br60, Ffm, and Shishito_GE).

Among the four newly isolated phages (Reaper_GE, Savina_GE, Aji_GE and Shishito_GE), Reaper_GE showed strict requirements for O-antigen including a complete LPS (Figure 2B), while Savina_GE primarily showed dependency on O-antigen followed by inner core mutants, and outer core mutants (Figure 2B, Dataset S4). For T5-like phage Aji_GE, both *fepA* (TonB-dependent enterobactin receptor) and *oafA* (O-antigen acyltransferase) showed high fitness scores. OafA performs an acetylation reaction on the abequose residue to create the O5-antigen serotype in LT2-derived strains (Slauch et al., 1996), and probably enhances infection via gaining access to the FepA-TonB complex. Related phenomena have been observed for other *Demerecviridae*, where other O-antigen modifications facilitated increased phage susceptibility (Heller and Braun, 1982; Kim and Ryu, 2012). Finally, similar to Br60 and Ffm phages isolated on rough-LPS *Salmonella*, Shishito_GE displayed strong host fitness defects in many of the LPS and O-antigen mutants in our library.

**Figure 2:**
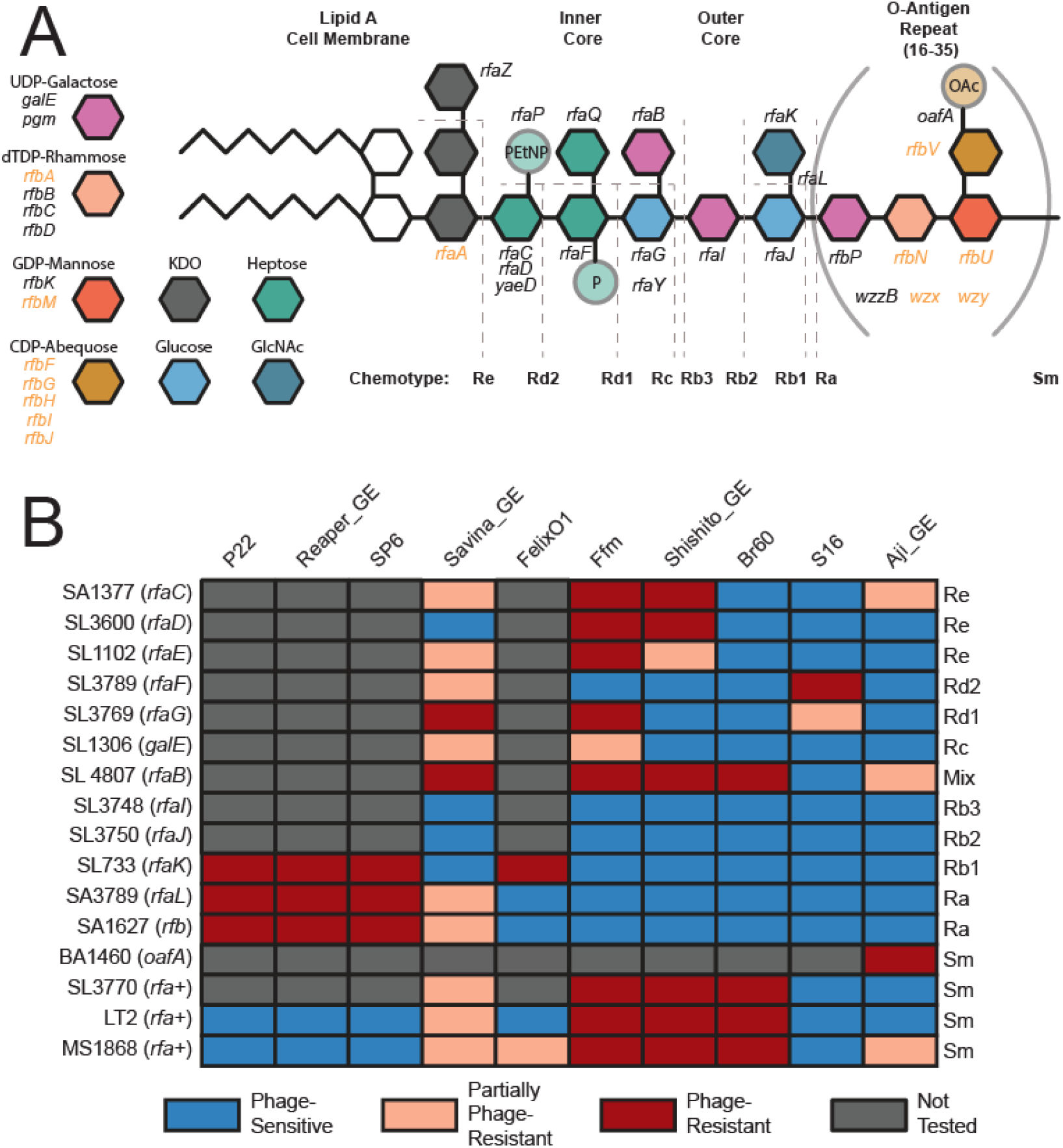
Validation of LPS-moiety requirements for *Salmonella* phages. (A) Overview of O5 *S*. Typhimurium LPS and O-antigen biosynthesis. The four sugars in brackets comprise the O-antigen, which repeats 16-35 times per LPS molecule under standard growth conditions. Key for non-essential LPS and O-antigen precursor biosynthesis genes are described to the right. Genes covered in our library and used for analysis are written in black. Genes not covered in our library, and thus not analyzed in this study are written in orange. (B) Infectivity matrix using a previously established *Salmonella* LPS panel (Table S5). The identity of the LPS chemotype corresponding to specific mutation is presented in (A). sm stands for smooth-LPS chemotype. Data for this figure is aggregated from Figures S5-S11, S13-S15, and S17.

To validate some of the top phage resistance phenotypes from our genetic screens, we used an established collection of *Salmonella* mutant strains in addition to the construction of deletion strains (Table S5)(Roantree et al., 1977; Sanderson et al., 1974). To confirm the role of OafA and TonB-dependent enterobactin receptor FepA (FepA-TonB complex) on Aji_GE infectivity, we constructed strains with deletions in *oafA, fepA*, and *tonB* (Methods). Aji_GE phage plaque assays on these strains confirmed the essentiality of OafA and both FepA-TonB in infection (Figures 2 and S17). For phages that showed stringent requirements of O-antigen and LPS, we used an established chemotype-defined LPS mutant panel in a *S*. Typhimurium strain background that is closely related to our LT2 derivative (Figures 2, S5-S11, and S13-S15, Table S5, Methods). Our phage infectivity results on the LPS chemotype panel are in agreement with earlier published data for some of the phages used in this work (Bohm et al., 2018; Lindberg and Hellerqvist, 1971; Marti et al., 2013; Mutalik et al., 2020; Wright et al., 1980) and consistent with our high-throughput genetic screens for all phages (Figure 1B). For example, LPS chemotype panel data confirmed the strict requirements for O-antigen including a complete LPS for Reaper_GE infectivity. We confirmed that Savina_GE most efficiently infects strains with an incomplete outer core, but less so against strains without O-antigen or strains missing outer core entirely (Figures 2 and S9). This result indicates Savina_GE preferentially employs LPS as a receptor, but branched LPS residues such as those added by *rfaK* and O-antigen biosynthesis probably hinder efficient adsorption. Though OafA activity is important for Aji_GE infection (Figure 2, BA1460), the acetylation provided by OafA activity does not seem to be critical in the absence of complete LPS and O-antigen as those mutants showed significant infection (Figures 2, S3, S11, and S17). The plaque assays of Shishito_GE on the LPS mutant panel confirmed that, like phages Br60 and Ffm, it only infects rough-LPS strains of *Salmonella* (Figures 2, S4, and S13-S15). As the inner and outer core LPS structure of *S*. Typhimurium is conserved in *E. coli* K-12, we confirmed these observations using data from an RB-TnSeq library of *E. coli* K-12 (Figure S4, Text S1, Methods). In summary, the combination of our high-throughput genetic screen and assays on single-gene deletion strains provided higher resolution mapping of O-antigen, LPS or protein receptor requirement for all 11 phages in *Salmonella* (a detailed description for each phage is in Text S1).

### Discovery of Novel Cross-Resistant Genotypes Between Diverse Phages

Next, we looked at the number and pattern of high-fitness scoring genes against our panel of phages to identify similarity in infectivity cycles and commonality in genetic barriers leading to phage cross-resistance. The most studied mode of resistance between phages is when they share a common receptor (for example, phages binding to LPS), and any modification in the common receptor yields cross-resistance to those phages (Chan and Turner, 2020; Mutalik et al., 2020; Shin et al., 2012; Wright et al., 2018, 2019). Though it is possible that other host factors are important for the infectivity cycle of different phages and can impart phage cross-resistance phenotypes, it remains a challenge to identify such non-receptor host factors and their role in phage infection. Thus they are not widely reported, nonetheless in the context of phage cross-resistance. For example, mutations in global transcriptional regulators can impart broad resistance to diverse phages that bind to different receptors, but have been proposed to impart higher fitness costs which probably explain their lower frequency of emergence(Betts et al., 2016; Díaz-Muñoz and Koskella, 2014; Hesse et al., 2020; Mutalik et al., 2020; Wright et al., 2018, 2019).

To gain more insights into phage cross-resistance, we compared the genes that show high-fitness scores across the 8 smooth-LPS binding phages screened in our study (Figure 3AB). The pair-wise comparison between any two phages indicated that, there is a wide range of shared high-fitness scoring genes. As expected, phages that bind to the same receptor shared many high-scoring genes indicating potential cross-resistance between them. For example, P22 SP6 and Reaper_GE bind to O-antigen and share many common high scoring hits. Conversely, there are instances of no high-fitness scoring genes shared between phages employing different receptors (for example, between Aji_GE and the O-antigen requiring phages Reaper_GE, SP6 and Savina_GE) (Figure 3A). Unexpectedly, we also observed instances of shared genes across phages that bind to different receptors, and point to a role played by the non-receptor host factors (Figure 3). For example, Aji_GE and FelixO1 have different receptors, yet they share a large number of high-fitness scoring genes, indicating potential cross-resistance independent of their primary receptors (Figures 1 and 3). Out of 52 non-receptor genes conferring resistance to FelixO1 and 32 non-receptor genes conferring resistance to Aji_GE, 29 were common to Aji_GE and FelixO1. These common non-receptor host factors appear to play diverse roles, and the functions they encode include disruptions across central metabolism (*aceEF, pta, ackA, fabF*), amino acid biosynthesis and regulation (*rpoN, glnDLG, ptsIN, aroM*), global regulation (*himAD, crp, rpoN, lon, arcB*), ion transport (*trkAH*), peptide transport (*sapABCF*), secondary messenger signaling (*gppA, cyaA*), translation (*trpS*), and other genes with less clear functions (*nfuA, yfgL, ytfP*). Some of these genes were recently implicated as host-factors in phage resistance in related organisms (Cowley et al., 2018; Goosen and Putte, 1995; Wright et al., 2018), though their role in phage cross-resistance and mechanisms were not determined. For example, *trk, sap, ace* and *rpoN* were recently associated with diverse phage resistance in *E. coli* (Cowley et al., 2018; Kortright et al., 2020) and *P. aeruginosa* (Wright et al., 2018). *himA* and *himD* (i.e. integration host factor subunits alpha and beta) are known to be involved in temperate phage infection pathways, though not shown for obligately lytic phages such as Aji_GE and FelixO1(Goosen and Putte, 1995).

**Figure 3:**
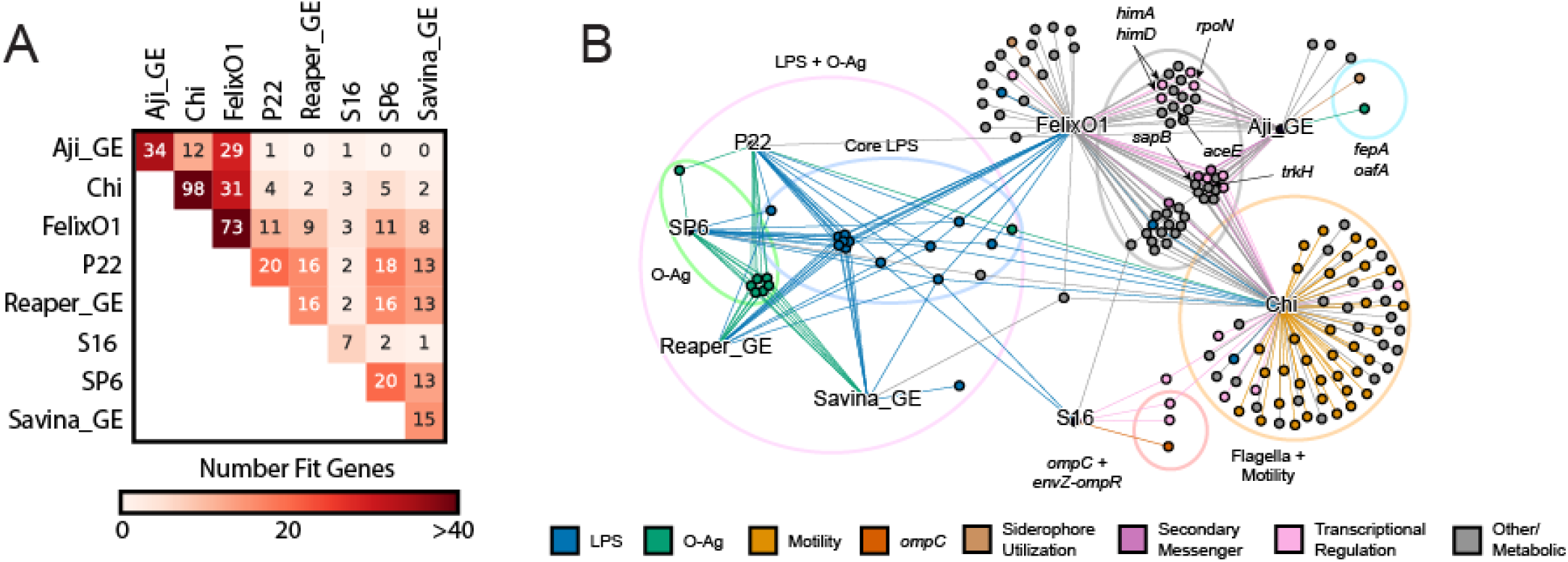
Cross-resistance is common between *Salmonella* phages. (A) Summary of cross-resistance patterns between phages observed in our screens. Heatmap color represents the total number of shared gene disruptions in *S*.Typhimurium that yield resistance to both phages. (B) Mixed-node network graph showing connections between phage nodes (text labels, black) and gene nodes (colored nodes). A gene node is connected to a phage node if disruptions in that gene gave high fitness against that phage (Dataset S4). Gene nodes are colored by encoded function. Notable gene function groupings and genes are additionally highlighted. Figures 3A and 3B are created from Dataset S4 using Supplemental Code - Figure3AB.

To investigate if these mutants indeed display cross-resistance to both Aji_GE and FelixO1, we selected a few top scoring genes to study further: *trkH, sapB, aceE, rpoN, himA*, and *himD*. For each of these 6 genes, we created individual mutants (Methods) and assessed Aji_GE and FelixO1 phage infectivity. Indeed, the *trkH, sapB, aceE, rpoN, himA* and *himD* mutants showed increased resistance to both FelixO1 and Aji_GE (Figures S19-S20). Consistent with prior reports of high fitness costs being associated with non-receptor phage cross-resistant mutants (Wright et al., 2018), mutants in *aceE, rpoN*, and *himA* displayed significant growth defects during planktonic growth, but were sufficiently fit to be uncovered in our screens. Some of these genes (for example, potassium transporter Trk and nitrogen assimilation sigma factor RpoN) are known to play an important ecological role in *Salmonella* virulence and fitness in infection contexts (Klose and Mekalanos, 1997; Su et al., 2009), indicating these phage resistance loci may exhibit an evolutionary trade-off with virulence.

### Sigma Factor Interplay Mediates Phage Cross-Resistance in *Salmonella*

To better identify the genetic basis of the phage cross-resistance phenotype imparted by *trkH, sapB, rpoN*, and *himA* mutants, we carried out RNA-Seq experiments and investigated whole-genome expression-level differences for each deletion compared to wild-type MS1868 (N=3 for all except for *himA*, which was N=2). In aggregate, we observed 635 differentially expressed genes (among which 437 are unique to one of the knock-out strains) in *trkH, sapB, rpoN*, and *himA* mutants compared to wild-type (Figure 4A, Dataset S6). To the best of our knowledge, none of the differentially expressed genes were related to FelixO1’s and Aji_GE’s suspected receptors (LPS and FepA respectively). In addition, neither of the known innate immunity defense mechanisms in *S*. Typhimurium (type I CRISPR or type I BREX), were found to be differentially expressed in any of these genetic backgrounds (Barrangou and Oost, 2014; Shariat et al., 2015). Thus we suspected this mode of resistance was likely due to global regulatory changes. We focused our analysis to *trkH, sapB*, and *rpoN* mutant backgrounds that showed upregulation of the *spv* virulence operon (*spvABC*), located on the PSLT plasmid native to *S*. Typhimurium (Dataset S6). In addition to being studied for its essentiality in *Salmonella* virulence, the *spv* operon is also well-known for being regulated by RpoS, a general stress response sigma factor (Chen et al., 1995; Fang et al., 1992; Nickerson and Curtiss, 1997). As the RpoS regulation is well-studied in *E. coli* and *S*. Typhimurium (Battesti et al., 2011; Hengge-Aronis, 2002; Ibanez-Ruiz et al., 2000; Lago et al., 2017; Lévi-Meyrueis et al., 2014; Lucchini et al., 2009; Nickerson and Curtiss, 1997), we looked for expression changes in RpoS-dependent genes in *trkH, sapB*, and *rpoN* mutant backgrounds. We found that a number of known RpoS-regulated genes were significantly upregulated versus wild-type (passing thresholds of log2FC > 2, p_adj < 0.001) (Figure 4CDE, Dataset S6), further implicating RpoS involvement in resistance to both Aji_GE and FelixO1 phages (Figure 4B).

**Figure 4:**
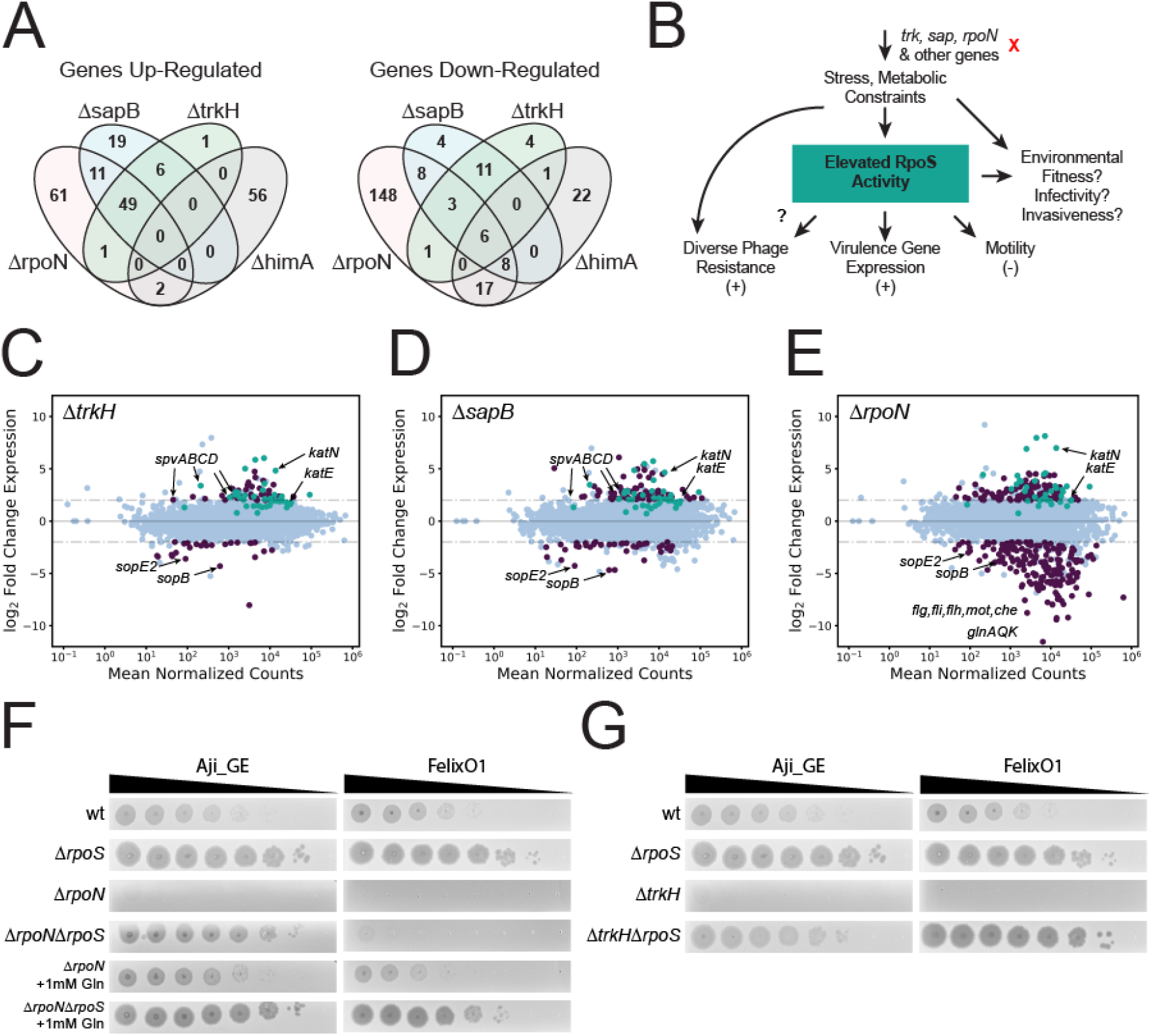
Cross-resistance mechanisms are mediated by RpoS. (A) Summary of genes with significant up- and down-regulation relative to wild-type for *sapB, trkH, rpoN*, and *himA* mutants. Reported values are genes with log2-fold changes over 2 and Bonferroni-corrected p values below 0.001. (B) Proposed model for phage cross-resistance observed in this study. Loss of function of genes such as *trkH, sapB*, or *rpoN* impose stress and metabolic constraints on *S*. Typhimurium. In some cases, this elevates RpoS activity and leads to multi-phage resistance. However, the environmental fitness, virulence, and invasiveness implications of these mutants are not known. (CDE) MA-plots for differential expression data for (C) MS1868Δ*trkH*, (D) MS1868ΔsapB, and (E) MS1868Δ*rpoN* mutants over wild-type MS1868. Differentially expressed genes (abs(log2FC ≥2), Bonferroni-corrected p values below 0.001) are shown in purple. RpoS-regulated genes are shown in teal based on a curated list from (Lucchini et al., 2009). Specific genes are highlighted for emphasis including RpoS-activity indicators *katE* and *katN*. (F) Aji_GE and FelixO1 phage susceptibility assays focused on Δ*rpoN*-mediated phage resistance. For both phages supplementing with glutamine (Gln) restores phage infectivity in Δ*rpoN* context. A secondary deletion in *rpoS* is sufficient to restore Aji_GE infectivity in a Δ*rpoN* strain. However, FelixO1 is only restored with additional supplementation of glutamine. (G) Aji_GE and FelixO1 phage susceptibility assays focused on Δ*trkH*-mediated phage resistance. A secondary deletion in *rpoS* is sufficient to restore both Aji_GE and FelixO1 infectivity in a Δ*trkH* strain. Figures 4ACDE are created from Dataset S6 using Supplementary Code - Figure4.

The general stress response sigma-factor RpoS activity in *Salmonella* is critical for many aspects of its adaptive lifestyle, including general virulence (Lévi-Meyrueis et al., 2014; Lucchini et al., 2009). However, comparative studies in clinical isolates of *Salmonella* found decreased RpoS activity in model strain LT2 versus related virulent strains due to a suboptimal start codon (Wilmes-Riesenberg et al., 1997). As a LT2 derivative, MS1868 has this suboptimal codon (Graña et al., 1985), so it is intriguing to find signatures of elevated RpoS activity and virulence-associated *spv* expression in phage resistant candidates. To confirm the impact of RpoS on phage infection, we created a *rpoS* deletion mutant and additional double gene replacement mutants of *rpoS* with one of *trkH, sapB*, or *rpoN*. The single *rpoS* deletion mutant displayed increased sensitivity to both FelixO1 and Aji_GE phage (Figures 4FG and S19-S20). In addition, the *rpoS* deletion was also sufficient to restore infectivity in *trkH, sapB*, and *rpoN* mutants to levels observed in *rpoS* mutants (Figures 4FG and S19-S20). While *himA* mutants did not show elevated levels of RpoS activity in our RNA-Seq data, we suspect that phage-resistance in many mutants within the Aji_GE and FelixO1 cross-resistance network emerged from RpoS activity beyond these mutants. More broadly, RpoS activity likely plays a role in intermediate phage-resistance phenotypes that are typically difficult to quantify in pooled fitness assays, but observable for these two phages.

In the *rpoN* (encoding sigma factor-54) mutant background, the *rpoS* mutation was sufficient to restore infectivity of phage Aji_GE, but insufficient to restore infectivity of FelixO1 (Figures 4F and S19-S20). Like RpoS, the alternate sigma factor RpoN is known to regulate a diverse set of pathways involved in adaptation and survival in unfavorable environmental conditions including nitrogen starvation. Because *rpoN* mutants decrease glutamine uptake and biosynthesis and have significant growth defects, the phage resistance phenotype observed in *rpoN* mutants potentially indicate the importance of glutamine levels on successful phage infection (Dataset S4) (see also: (Aurass et al., 2018; Samuels et al., 2013)). To assess the dependence of glutamine on phage resistance mechanism, we repeated phage infection supplemented with glutamine in *rpoN* mutants. Both FelixO1 and Aji_GE were able to successfully plaque on *rpoN* mutants supplemented with glutamine. In the *rpoN, rpoS* double mutant background, additional glutamine supplementation was able to nearly restore FelixO1 infectivity to the *rpoS* mutant’s baseline (Figures 4F and S19-S20). Thus, we propose *rpoN* loss-of-function probably manifests two avenues of phage resistance. First, nutrient limitation to the cell can “starve” phage replication, such as FelixO1 but not Aji_GE, during infection. Second, elevated RpoS activity (likely induced by nutrient limitation) confers further resistance to phage infection, extending to diverse phages such as FelixO1 and Aji_GE. In summary, these studies uncover intricate interplay between host factors and nutritional status of the cell in phage cross-resistance phenotype.

### Investigation into phage sensitivity of natural *Salmonella* strain variants

Finally, we wondered how gene requirements uncovered in our genome-wide genetic screens corresponded to naturally occurring variation in and phage sensitivity of *S*.Typhimurium isolates. More broadly, we were interested in to what degree these gene requirements in a model strain were predictive of phage sensitivity patterns in closely related strains. Though phage host range determination using a panel of strains belonging to a species of bacterium is a century old practice, the genetic basis of the phage infectivity pattern has remained unresolved(Holmfeldt et al., 2007; Hyman and Abedon, 2010; de Jonge et al., 2019; Moller et al., 2019; Weitz et al., 2013). For example, phage infectivity patterns using a panel of phages (phage typing) to discriminate *Salmonella* serovars for epidemiological investigation/surveillance is even practiced today, while the infectivity pattern is not typically investigated mechanistically(Chirakadze et al., 2009; Rabsch, 2007). We hypothesized that the similarity and differences in genetic determinants involved in phage resistance might be able to explain the genetic basis of phage infectivity when extended to a panel of *Salmonella* strains. To assess the relationship between genomic content and phage sensitivity among natural strain variants, we sourced a panel of 21 *S*. Typhimurium strains belonging to the SARA collection(Beltran et al., 1991). We also included a model nontyphoid clinical isolate D23580 from Malawi(Canals et al., 2019) and ST4/74 strain, originally isolated from a calf with salmonellosis(Richardson et al., 2011) as a reference. The SARA collection is a set of strains of *Salmonella* isolated from a variety of hosts and environmental sources in diverse geographic locations, classified into 17 electrophoretic types, observed variation in natural populations and is reflective of much of the diversity identified in panels derived from recent *S*. Typhimurium outbreaks(Fu et al., 2015).

We re-sequenced these 21 strains to confirm their identity and assembled their genomes as described in Methods. Our analysis showed that all isolates, except for SARA7 and SARA8, have a close phylogenetic relationship (>99% pairwise average nucleotide identity, ANI) in agreement with an earlier report(Fu et al., 2015). Next, we searched for the 184 unique high-scoring gene hits uncovered in this work (Table S4) across our panel of *Salmonella* genomes and observed little variation in the sequence of genes, though there might be changes in expression and activity (Dataset S8). Among the key differences in our gene content analysis, we observed nonsense mutations or frame shifting changes in the coding region of *rfaK* in SARA20, *ompC* and *rfbN* in SARA6 and *oafA* in SARA9 compared to our reference strain *Salmonella* LT2. Mutation in the coding region of *rfaK/waaK* in SARA20 yields two truncated proteins, and neither of them have a complete glycosyltransferase domain. It is known that *rfaK* mutants lack the GlcNAc residue in the LPS outer core and are also unable to express O-antigen because this GlcNAc residue is essential for the recognition of core oligosaccharide acceptor by the O-antigen ligase WaaL(Hoare et al., 2006). We postulated that absence of outer core GlcNAc (the biosynthetic product of RfaK) in SARA20 probably alters the structure of O-antigen and may yield resistance to O-antigen binding by P22, SP6, Reaper_GE and FelixO1 phages. Disruption in the *ompC* coding region in SARA6 may compromise S16 phage infectivity and disruption in *oafA* coding region (in SARA9) probably interferes with efficient infection by Aji_GE. Broadly, our analysis predicts that all 23 *Salmonella* isolates except the ones mentioned above should show similar phage infectivity patterns as compared to the laboratory strain used in our genetic screens.

To assess the infectivity pattern of the 11 phages against the 23 *Salmonella* strains, we carried out standard spotting assays. Figure 5 shows the phage infectivity data and phylogenetic distance between *Salmonella* strains, with a phylogenetic tree built from gene sequences of 115 single-copy marker genes (Methods). In agreement with our genome-based prediction, 22 strains (out of 23 strains) displayed broad sensitivity to all O-antigen binding phages (except Ffm, Br60 and Shashito-GE) (Figures 1-2 and 5). SARA6 was the only strain sensitive to Ffm, Br60 and Shashito_GE phages and was also resistant to all phages binding O-antigen (P22, SP6, Reaper_GE), indicating SARA6 may have rough-LPS phenotype. Analysis of SARA6 genome indicated that *rfbN*, a gene encoding rhamnosyltransferase important for O-antigen synthesis has a mutation and that this strain would not be able to express O-antigen, in agreement with its resistance to O-antigen binding phages (P22, SP6 and Reaper_GE) while showing sensitivity to core LPS binding phages (Ffm, Br60 and Shashito-GE). Disruption of the *ompC* coding region in SARA6 while retaining infectivity with S16 phage indicates there is a possibility of OmpC independent infectivity pathway as seen in some T4-like phages(Washizaki et al., 2016). SARA20 showed no sensitivity to both smooth-LPS binding (P22, SP6, Reaper_GE, FelixO1) and rough-LPS binding phages (Ffm, Br60 and Shashito-GE), raising an interesting question about its LPS architecture. Though mutation in the coding region of *rfaK* and absence of O-antigen in SARA20 explains its resistance to P22, SP6, Reaper_GE, FelixO1 phages, though the resistance showed by rough-LPS binding phages Ffm, Br60 and Shashito-GE indicate additional factors likely play a role. Finally, in agreement with our gene content analysis (above), SARA9, with disruption in the *oafA* coding region, showed inefficient infection by Aji_GE.

**Figure 5.**
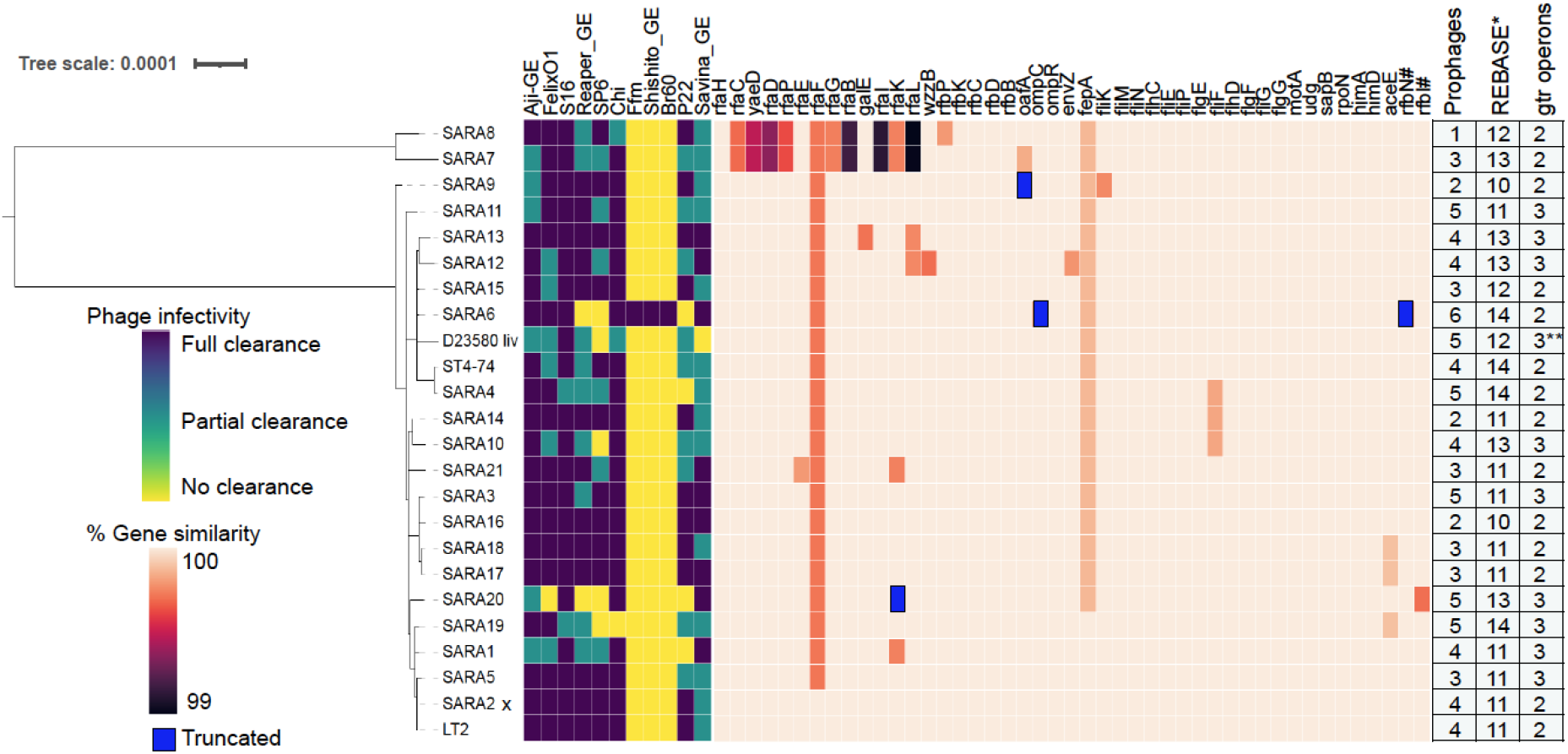
Host range of *Salmonella* phages and conservation of host factors involved in phage infection. Infectivity pattern of 11 phages on a panel of 23 *Salmonella* strains was inferred by spotting assay. Phylogenetic relationships of the *Salmonella* strains were estimated by phylogenetic analysis of 115 single-copy marker genes (Methods). Gene similarity was calculated by TBLASTN search with LT2 proteins in SARA genomes for 184 unique gene hits uncovered in this work, and 45 (out of 184) are shown in this figure (Supplementary Datasets S4 and S8) with total number of predicted prophages, restriction/modification proteins and *gtrABC* operons. Gens with premature stop codons/truncations due to insertion or deletion (compared to LT2) are marked blue. Complete detail of gene similarity of 184 unique gene hits, prophages, restriction/modification proteins and *gtrABC* operons across 23 *Salmonella* strains are given in Datasets S8 and S9 * pseudogenes and incomplete genes were excluded from the analysis, SARA2 x indicates LT2 strain. ** BTP1 prophage of D23580 contains only *gtrA* and *gtrC* genes; # denotes genes that are not from the genetic screen. Figure 5 was created from Datasets S8 and S9.

In addition to these broad agreements between gene content analysis and phage infectivity, the genetic basis of strong phage resistance showed by SARA1 and SARA4 (for P22), SARA10 (for SP6), D23580 (for SP6 and Savina_GE) and SARA19 (for SP6, Chi) is unclear. We also observed a partial clearance pattern in our spot tests for many phages across 23 isolates, probably indicating inefficient infection cycles or partial inhibition characterized by turbid plaques. Overall, these results indicate that phage host range is probably defined by additional constraints than the host factors uncovered in our genetic screens. To look for other host factors that might be playing a role in efficient phage infection, we bioinformatically searched for prophages, genes encoding O-antigen modification systems (*gtrABC* operon) and restriction modification systems encoded in our panel of 23 *Salmonella* strains. The role played by these genetic elements on phage infectivity and resistance are well appreciated(Bernheim and Sorek, 2020; Bondy-Denomy et al., 2016; Cota et al., 2015; Davies et al., 2013; Dedrick et al., 2017; Dy et al., 2014; van Houte et al., 2016; Owen et al., 2020; Rostøl and Marraffini, 2019; Samson et al., 2013; Vasu and Nagaraja, 2013; Wahl et al., 2019). Our comparative analysis provided a list of strain-specific restriction/modification and O-antigen modification genes that may influence phage infection outcome. However, we could not identify a single genomic loci, whose presence or absence fully coincides with the strong phage resistance in the SARA1, SARA4, SARA10, SARA19 and D23580 strains in addition to the inefficient phage infectivity pattern across our panel of *Salmonella* strains. We postulate that a combination of genetic factors rather than a single gene mutation probably drive the smaller changes in phage infective efficiency (Datasets S8 and S9).

## Discussion

Here, we employed an unbiased RB-TnSeq loss-of-function approach to uncover the genetic determinants important in phage infection and resistance in a model enteric *Salmonella* species across 11 distinct dsDNA phages, including four phages from a therapeutic formulation. In addition to identifying known receptors for model *Salmonella* phages, our genome-wide screens identify novel receptor and non-phage-receptor host factors important for a panel of dsDNA phages. We validate many of these high fitness hits via single gene deletion strains. Our results indicate diverse modes of phage resistance including disruption in the phage infectivity pathway downstream from phage receptors. Characterization of these non-receptor phage resistance factors shared between two unrelated phages (indicating cross-resistance) identified an intricate interplay between alternative sigma factors pointing to how phage predation might be influenced by growth and nutritional status of the cell. Finally, our host-range investigation of 11 phages across a panel of 23 *Salmonella* strains showed differences in the infectivity pattern of some phages despite having high conservation of top scoring hits in our genome-wide screens in a closely related model organism. Comparative analysis identified instances where sequence variation in target receptor explained some of the phage susceptibility pattern, but there are additional factors and interactions both in the target host and phages that defines the host range. Overall this study highlights the importance of unbiased high-throughput genetic screens across a panel of phages in uncovering diversity of host factors important phage infection, provides insights on the genetic basis and modes of cross-resistance between sets of phages, and uncover gaps in our understanding of phage host-range across natural bacterial isolates.

Our genome-wide screens also suggest how phage selection can be used to drive beneficial tradeoffs to modulate pathogen virulence, sensitivity and fitness. For example, LPS and O-antigen play a critical role in the lifestyle of *Salmonella* virulence and have a myriad of effects on phage predation. Phage selection to drive truncation, loss, or reduction of LPS and O-antigen in *Salmonella* could be employed to decrease virulence and increase its susceptibility to antibiotics, decreased swarming motility, decreased colonization, and decreased fitness (Kong et al., 2011; Nagy et al., 2006; Toguchi et al., 2000). Our investigation into a network of cross-resistant genotypes against unrelated phages FelixO1 and Aji_GE led to the role of RpoS and RpoN activity, a virulence-regulating alternative sigma factors in *Salmonella sp* (Chen et al., 1995; Fang et al., 1992; Wilmes-Riesenberg et al., 1997) on phage infectivity. Specifically, the association of increased RpoS activity with phage resistance raises intriguing ecological questions for consideration. While RpoS activity is associated with virulence, are these phage-resistant genotypes more virulent and fit in infection contexts? Further, do these genotypes display increased RpoS activity and/or virulence in more virulent *Salmonella* strains? If so, the selection for increased phage-resistant strains with increased virulence-associated RpoS activity would be a deleterious outcome from therapeutic phage predation and an undesirable criterion for potentially therapeutic phages. Conversely, does the lack of phage predation contribute to the neutral drift of RpoS alleles and virulence in laboratory settings (Robbe-Saule et al., 1995; Wilmes-Riesenberg et al., 1997)? It is known that RpoS directly and indirectly regulates more than 10% of all genes in *E. coli* (Battesti et al., 2011) and *S*. Typhimurium (Lago et al., 2017; Lévi-Meyrueis et al., 2014), and is involved in adaptation to diverse environments and metabolic states (Battesti et al., 2011, 2015; Hengge-Aronis, 2002; Lucchini et al., 2009; Nickerson and Curtiss, 1997; Robbe-Saule et al., 1995). Thus, phage resistance phenotypes associated with RpoS activity may be acting through activation of RpoS-mediated stress response pathways rather than the direct loss of RpoS itself. In some cases, dual regulation by genetic or nutritional factors and RpoS could lead to compensation. Some of these genotypes (for example,mutants in *trk, sap, ace*, and *rpoN*) were recently associated with phage resistance in *E. coli* (Cowley et al., 2018; Kortright et al., 2020) and *P. aeruginosa* (Wright et al., 2018), but are not yet linked to RpoS activity. Future work will explore if we see similar dependencies of alternative sigma-factors on phage resistance phenotypes in other pathogens (Dong and Schellhorn, 2010).

Our investigation into the host-range of *Salmonella* phages across closely related *Salmonella* isolates indicated that the highly fit phage resistance genotypes uncovered via genome-wide genetic screens are not complete predictors of phage infectivity and host range. Though these host factors showed little variation in their sequences across our panel of *Salmonella* isolates, it is possible that they vary in expression and activity, sufficient to impact phage infectivity cycle. The host-range of phages is not only defined by whether the host is susceptible to phage infection, but also on how phages evade host defences and overcome barriers to efficient infection. For example, Because O-antigen structures in *Salmonella typhimurium* sp. can comprise over 400 sugars per O-antigen LPS molecule(Crawford et al., 2012), it is no surprise that many bacteriophages adsorb to this highly exposed structure. However, many bacteriophages that adsorb to centralized outer membrane receptors can be occluded from their native receptor by the O-antigen structure (Domínguez-Medina et al., 2020). Systematic studies exploring the form and structure of O-antigens and how they impact accessibility of phage receptors are needed. We postulate that there are likely additional constraints affecting optimal phage infectivity, and probably we might have missed uncovering these additional factors in our model strain because of highly fit phage receptor mutants. Considering differences in phage infectivity in a panel of strains with highly conserved genetic determinants, we posit that systematic study of differences in transcriptional and translation processes in these strains might provide more insights as illustrated in a few recent phage-host interaction studies(Brandão et al., 2021; Howard-Varona et al., 2018). Future studies could also employ recently developed methods(Mutalik et al., 2019; Rishi et al., 2020; Thibault et al., 2019) to provide higher resolution into phage-host interactions and may aid in filling the knowledge gaps on phage host-range. These methods could be extended to a few closely related and phylogenetically distant strains to understand the variability in host factors impacting phage infectivity patterns. Finally, by combining the genetic tools developed for functional assessment of host genes with targeted or genome-wide loss-of-function mutant libraries in few model phages, can provide additional insights into the host specificity of phages.

As high-throughput genetic screens to understand phage-host interactions grow more commonplace across diverse bacteria (Bohm et al., 2018; Christen et al., 2016; Cowley et al., 2018; Mutalik et al., 2019; Pickard et al., 2013; Price et al., 2018; Rishi et al., 2020; Rousset et al., 2018; Wetmore et al., 2015), leveraging fitness data across phages and bacterial genetic diversity constitutes a major challenge and opportunity. Further screens against antibiotics, such as those presented in earlier (Price et al., 2018), could rapidly discover collateral sensitivity patterns wherein phage resistant genotypes display sensitization to antibiotics or ecologically-relevant conditions (for instance sera or bile salts). Such information has the potential to form the basis of successful combinations of treatments (Chan et al., 2018; Kortright et al., 2019; Mangalea and Duerkop, 2020). We posit that phage-host interaction studies across diverse bacterial isolates in a range of biotic and abiotic conditions powered with novel transcriptomics and proteomics tools can provide rich datasets for host-range predictive models and rational phage cocktails formulations.

## Materials and Methods

### Bacterial strains and growth conditions

Strains, primers, and plasmids are listed in Tables S3-S5, respectively. *S*. Typhimurium LT2 derivative strain MS1868 genotype is *S*. Typhimurium LT2 (leuA414(Am) Fels2-hsdSB(r-m+))(Graña et al., 1985). In general, all *Salmonella* strains were grown in Luria-Bertani (LB-Lennox) broth (Sigma) at 37°C, 180 rpm unless stated otherwise. When appropriate, 50 µg/mL kanamycin sulfate and/or 34 µg/mL chloramphenicol were supplemented to media. For strains containing an ampR selection marker, carbenicillin was employed at 100 µg/mL, but exclusively used during isolation of clonal mutants to avoid mucoidy phenotypes. All bacterial strains were stored at −80°C for long term storage in 25% sterile glycerol (Sigma).

### Bacteriophages and propagation

Bacteriophages employed in this study and sources are listed in Table 1. All phages were either successively serially diluted or streaked onto 0.7% LB-agar overlays for isolation. For bacteriophage Chi, 0.35% LB-agar overlays were employed. Bacteriophage Aji_GE_EIP16, Reaper_GE_8C2, Savina_GE_6H2, and Shishito_GE_6F2 were isolated from a commercial bacteriophage formulation from Georgia. All phages isolated from this source are denoted with “_GE” (to recognize being sourced from Georgia). All other bacteriophages were re-isolated from lysates provided from stock centers or gifts from other labs (Table 1). Bacteriophage Aji_GE_EIP16, Chi, FelixO1, P22 (a strictly lytic mutant), Reaper_GE_8C2, S16, and SP6 were isolated and scaled on *S*. Typhimurium MS1868. Bacteriophage Br60, Ffm, Savina_GE_6H2, and Shishito_GE_6F2 were isolated and rough-LPS mutant *S*. Typhimurium. SL733 (BA1256). We followed standard protocols for propagating phages (Kutter and Sulakvelidze, 2004). Br60, Chi, Ffm, P22, Reaper_GE_8C2, S16, Savina_GE_6H2, Shishito_GE_6F2, and SP6 were propagated in LB-Lennox liquid culture on their respective strains. Reaper_GE_8C2, Savina_GE_6H2, and Shishito_GE_6F2 were additionally buffer-exchanged into SM-Buffer (Teknova) via ultrafiltration (Amicron 15) and resuspension. Bacteriophage Aji_GE_EIP16 and FelixO1 were propagated in LB-Lennox liquid culture on their respective strains and further propagated through a standard overlay method. Whenever applicable, we used SM buffer without added salts (Tekova) as a phage resuspension or dilution buffer and routinely stored phages as filter-sterilized (0.22um) lysates at 4°C.

Bacteriophages Aji_GE_EIP16, Reaper_GE_8C2, Savina_GE_6H2, and Shishito_GE_6F2 were additionally whole-genome sequenced and assembled. Approximately 1e9 PFU of phage lysate was gDNA extracted through Phage DNA Isolation Kit (Norgen, 46800) as per manufacturer’s instructions. Library preparation was performed by the Functional Genomics Laboratory (FGL), a QB3-Berkeley Core Research Facility at UC Berkeley. Sequencing was performed at the Vincent Coates Sequencing Center, a QB3-Berkeley Core Research Facility at UC Berkeley on a MiSeq using 75PE runs for Reaper_GE_8C2, Savina_GE_6H2, and Shishito_GE_6F2 and using 150SR run for Aji_GE_EIP16. Phage genomes were assembled using KBase (Arkin et al., 2018). Illumina reads were trimmed using Trimmomatic v0.36 (Bolger et al., 2014) and assessed for quality using FASTQC. Trimmed reads for Aji_GE_EIP16, Reaper_GE_8C2, and Shishito_GE_6F2 were assembled using Spades v3.13.0 (Nurk et al., 2013). Trimmed reads for Savina_GE_6H2 were assembled using Velvet v1.2.10 (Zerbino and Birney, 2008). The primary, high coverage contig from these assemblies was investigated and corrected for incorrect terminus assembly using PhageTerm v1.011 on CPT Galaxy (Garneau et al., 2017). In this manuscript, we limited analyses of these sequences to assessing phylogeny of these phages, which we performed with BLASTN (Dataset S7). A detailed genomic characterization will be published by Dr. Elizabeth Kutter. Sequences and preliminary annotations can be found at JGI IMG under analysis projects Ga0451357, Ga0451371, Ga0451358, and Ga0451372.

### Construction of MS1868 RB-TnSeq library

We created the *Salmonella enterica* serovar Typhimurium MS1868 (MS1868_ML3) transposon mutant library by conjugating with *E. coli* WM3064 harboring pHLL250 mariner transposon vector library (strain AMD290) (Figure S1). To construct pHLL250, we used the magic pools approach we outlined previously(Liu et al., 2018). Briefly, pHLL250 was assembled via Golden Gate assembly using BbsI from part vectors pHLL213, pHLL216, pHLL238, pHLL215, and pJW14(Liu et al., 2018). We then incorporated millions of DNA barcodes into pHLL250 with a second round of Golden Gate assembly using BsmBI. Briefly, we grew *S*. Typhimurium LT2 MS1868 at 30°C to mid-log-phase and combined equal cell numbers of *S*. Typhimurium LT2 MS1868 and donor strain AMD290, conjugated them for 5 hrs at 30°C on 0.45-µm nitrocellulose filters (Millipore) overlaid on LB agar plates containing diaminopimelic acid (DAP) (Sigma). The conjugation mixture was then resuspended in LB and plated on LB agar plates with 50 ug/ml kanamycin to select for mutants. After 1 day of growth at 30°C, we scraped the kanamycin-resistant colonies into 25 mL LB and processed them as detailed earlier to make multiple 1-mL − 80°C freezer stocks. To link random DNA barcodes to transposon insertion sites, we isolated the genomic DNA from cell pellets of the mutant libraries with the DNeasy kit (Qiagen) and followed published protocol to generate Illumina compatible sequencing libraries(Wetmore et al., 2015). We then performed single-end sequencing (150 bp) with the HiSeq 2500 system (Illumina). Mapping the transposon insertion locations and the identification of their associated DNA barcodes was performed as described previously (Price et al., 2018). In total, our 66,996 member pooled library consisted of transposon-mediated disruptions in 3,759 out of 4,610 genes, with an average of 14.8 disruptions per gene (median 12). Compared to a non-barcoded reported transposon mutant library in *S*. Typhimurium 14028s *(Porwollik et al*., *2014)*, we suspect 434 of the 851 unmutated genes are likely essential, and 380 likely nonessential. We abstain from interpreting essentiality of 37 additional genes due to inability to uniquely map insertions or due to gene content differences between the two libraries. Additional details for the composition of the *S*. Typhimurium MS1868 library can be found in Table S1 and Dataset S1.

### Liquid culture “competitive” fitness experiments

Competitive, phage-stress fitness experiments were performed in liquid culture, as phage progeny from an infection of one genotype could subsequently infect other host genotypes. All bacteriophages were tested against the MS1868 library. Bacteriophage Br60, Ffm, and Shishito_GE_6F2 were additionally tested against the previously described *E. coli* BW25113 library (Wetmore et al., 2015). To avoid jackpot effects, at least two replicate experiments were performed per phage-host library experiment as presented earlier (Mutalik et al., 2020). Briefly, a 1 mL aliquot of RB-TnSeq library was gently thawed and used to inoculate a 25 mL of LB supplemented with kanamycin. The library culture was allowed to grow to an OD600 of ∼1.0 at 37°C. From this culture we collected three, 1 mL pellets, comprising the ‘Time-0’ or reference samples in BarSeq analysis. The remaining cells were diluted to a starting OD600 of 0.04 in 2X LB with kanamycin. 350 µL of cells were mixed with 350 µL phage diluted in SM buffer to a predetermined MOI and transferred to a 48-well microplate (700 µL per well) (Greiner Bio-One #677102) covered with breathable film (Breathe-Easy). Phage infection progressed in Tecan Infinite F200 readers with orbital shaking and OD600 readings every 15 min for 3 hours at 37°C. At the end of the experiment, each well was collected as a pellet individually. All pellets were stored at −80°C until prepared for BarSeq.

### Solid agar “noncompetitive” fitness experiments

Noncompetitive, phage-stress fitness experiments were performed on solid-agar plate culture as presented earlier (Mutalik et al., 2020). Solid plate fitness experiments were performed by assaying all 11 bacteriophages against the MS1868 library. Bacteriophage Br60, Ffm, and Shishito_GE_6F2 were additionally assayed on the *E. coli* BW25113 library (Wetmore et al., 2015). For the solid plate experiments a 1 mL aliquot of the RB-TnSeq library was gently thawed and used to inoculate a 25 mL LB supplemented with kanamycin. The library culture was allowed to grow to an OD600 of ∼1.0 at 37°C. From this culture we collected three, 1 mL pellets, comprising the ‘Time-0’ for data processing in BarSeq analysis. The remaining cells were diluted to a starting OD600 of 0.01 in LB with kanamycin. 75 µL of cells were mixed with 75 µL of phage diluted in SM buffer to a predetermined MOI and allowed to adsorb for 10 minutes. The entire culture was spread evenly over a LB agar plate with kanamycin and grown overnight at 37°C. The next day, all resistant colonies were collected and suspended in 1.5 mL LB media before pelleting. All pellets were then stored at −80°C until prepared for BarSeq.

### BarSeq of RB-TnSeq pooled fitness assay samples

Genomic DNA was isolated from stored pellets of enriched and ‘Time 0’ RB-TnSeq samples using the DNeasy Blood and Tissue kit (Qiagen). We performed 98°C BarSeq PCR protocol as described previously (Mutalik et al., 2020; Wetmore et al., 2015). BarSeq PCR in a 50 uL total volume consisted of 20 umol of each primer and 150 to 200 ng of template genomic DNA. For the HiSeq4000 runs, we used an equimolar mixture of four common P1 oligos for BarSeq, with variable lengths of random bases at the start of the sequencing reactions (2–5 nucleotides). Equal volumes (5 uL) of the individual BarSeq PCRs were pooled, and 50 uL of the pooled PCR product was purified with the DNA Clean and Concentrator kit (Zymo Research). The final BarSeq library was eluted in 40 uL water. The BarSeq samples were sequenced on Illumina HiSeq4000 instruments with 50 SE runs. Typically, 96 BarSeq samples were sequenced per lane of HiSeq.

### Data processing and analysis of BarSeq reads

Fitness data for the RB-TnSeq library was analyzed as previously described (Wetmore et al., 2015). Briefly, the fitness value of each strain (an individual transposon mutant) is the normalized log2(strain barcode abundance at end of experiment/strain barcode abundance at start of experiment). The fitness value of each gene is the weighted average of the fitness of its strains. Further analysis of BarSeq data was carried out in Python3 and visualized employing matplotlib and seaborn packages. For heatmap visualizations, genes with under 25 BarSeq reads in the phage samples had their fitness values manually set to 0 to avoid artificially high fitness scores (due to the strong selection pressure imposed by phage predation).

Due to the strong selection pressure and subsequent fitness distribution skew resulting from phage infection, a couple additional heuristics were employed during analysis. Initially, per phage experiment, fitness scores were filtered for log2-fold-change thresholds, aggregated read counts, and t-like-statistics. Experiments using phages Ffm, Shishito_GE, and Br60 (which cannot infect wild-type MS1868, but can infect specific MS1868 mutants) against the MS1868 library employed negative thresholds to identify sensitized genotypes. A summary of log2-fold-change fitness and t-like statistic thresholds are provided in the Dataset S2. Each reported hit per phage was further processed via manual curation to minimize reporting of false-positive results due to the strong phage selection pressure. Here, all individual barcodes per genotype were investigated simultaneously for each experiment through both barcode-level fitness scores and raw read counts. First, genotypes were analyzed for likely polar effects. If the location and orientation of each fit barcode were exclusively against the orientation of transcription and/or exhibited strong fitness at the C-terminus of a gene, while being transcriptionally upstream of another fit gene, the genotype was likely a polar effect and eliminated. Second, genotypes were analyzed for jackpot fitness effects that could indicate a secondary site mutation. These cases were identified by investigating consistency between individual strains within a genotype. If the vast majority of reads per genotype belonged to a singular mutant (of multiple), we attributed the aggregate fitness score to secondary-site mutation effects and eliminated those genotypes from reported results. Genotypes where there were too few strains to make a judgment call on within genotype strain consistency (ie 1-3 barcodes) were generally excluded from analysis unless they were genotypes consistent with other high-scoring genotypes. Next, we investigated for consistency between read counts and fitness scores at both the strain level. In general, we found that strains with read counts under 25 often had inflated fitness scores under strong phage selection pressure and the subsequent fitness distribution skew resulting from phage infection. Cases where high fitness scores were attributed to a couple of strains with reads under 25 were eliminated as false positives as well. Finally, all genotypes were loosely curated for consistency across liquid experiments. Cases that barely passed confidence thresholds as described above that were inconsistent across replicate experiments were eliminated from reporting. A summary of fit genotypes that passed automated filtering and manual curation are reported in Dataset S4. No fit genotypes were added during manual analyses.

Network graphs were constructed using Gephi. Graph layout optimization was determined through a combination of manual placement of nodes (for instance phage nodes in Figure 3B) and layout optimization based off of equally weighted edges using the Yifan Hu algorithm. In all graphs, edges were calculated based on Dataset S4 using custom python scripts. In brief, in the mixed node graph in Figure 3B, edges were drawn with weight 1 between a phage node (fixed) and a gene node if that gene conferred resistance according to Dataset S4.

### Individual Mutant Creation

All individual deletion mutants in *S*. Typhimurium were created through lambda-red mediated genetic replacement (Sawitzke et al., 2007). Per deletion, primers were designed to PCR amplify either kanamycin or ampicillin selection markers with ∼30-40 bp of homology upstream and downstream of the targeted gene locus, leaving the native start and stop codons intact preserving directionality of gene expression at the native locus (Table S3). PCRs were generated and gel-purified through standard molecular biology techniques and stored at −20°C until use. All strains (including mutants) employed in this study are listed in Table S5.

Deletions were performed by incorporating the above dsDNA template into the *Salmonella* genome through standard pSIM5-mediated recombineering methods (Sawitzke et al., 2007). First temperature-sensitive recombineering vector, pSIM5, was introduced into the relevant *Salmonella* strain through standard electroporation protocols and grown with chloramphenicol at 30°C. Recombination was performed through electroporation with an adapted pSIM5 recombineering protocol. Post-recombination, clonal isolates were streaked onto plates without chloramphenicol at 37°C to cure the strain of pSIM5 vector, outgrown at 37°C and stored at − 80°C until use. For double deletions, this process was repeated two times in series with kanamycin followed by ampicillin selection markers. Gene replacements were verified by colony PCR followed by Sanger sequencing at the targeted locus (both loci if a double deletion mutant) and 16S rDNA regions (primers provided in Table S3).

### Assessing Phage Sensitivity

Phage-resistance and -sensitivity was assessed through efficiency of plating experiments. Bacterial hosts were grown overnight at 37°C. 100 µL of these overnight cultures were added to 5 mL of top-agar with appropriate antibiotics and allowed to solidify at room temperature. For assays including supplements such as glutamine, the supplement was added directly to the top agar layer. Phages were ten-fold serially diluted in SM Buffer, two microliters spotted out on the solidified lawn, and incubated the plates overnight at 37°C. Efficiency of plating was calculated as the ratio of the average effective titer on the tested host to the titer on the propagation host. For some assay strains, plaques showed diffused morphology and were difficult to count, or displayed plaque phenotypes distinct from its propagation host. In all cases, representative images are presented (Figures S5-S11, S13-S15). All plaquing experiments were performed with at least three biological replicates, each replicate occurring on a different day from a different overnight host culture.

### RNA-Seq experiments

Samples for RNA-Seq analysis were collected and analyzed for wild-type MS1868 (BA948) (N=3), knockout mutants for trkH (BA1124) (N=3) *sapB* (BA1136) (N=3), *rpoN* (BA1139) (N=3), and *himA* (BA1142) (N=2). All cultures for RNA-Seq were grown on the same day from unique overnights and subsequent outgrowths. Strains were diluted to OD600 ∼0.02 in 10 mL LB with appropriate selection marker, and then grown at 30°C at 180 RPM until they reached an OD600 0.4-0.6. Samples were collected as follows: 400 µL of culture was added to 800 µL RNAProtect (Qiagen), incubated for 5 minutes at room temperature, and centrifuged for 10 minutes at 5000xg. RNA was purified using RNeasy RNA isolation kit (Qiagen) and quantified and quality-assessed by Bioanalyzer. Library preparation was performed by the Functional Genomics Laboratory (FGL), a QB3-Berkeley Core Research Facility at UC Berkeley. Illumina Ribo-Zero rRNA Removal Kits were used to deplete ribosomal RNA. Subsequent library preparation steps of fragmentation, adapter ligation and cDNA synthesis were performed on the depleted RNA using the KAPA RNA HyperPrep kit (KK8540). Truncated universal stub adapters were used for ligation, and indexed primers were used during PCR amplification to complete the adapters and to enrich the libraries for adapter-ligated fragments. Samples were checked for quality on an Agilent Fragment Analyzer, but ribosome integrity numbers were ignored. This is routine for *Salmonella* sp., since they natively have spliced 23S rRNA (Burgin et al., 1990). Sequencing was performed at the Vincent Coates Sequencing Center, a QB3-Berkeley Core Research Facility at UC Berkeley on a HiSeq4000 using 100PE runs.

### RNA-Seq Data Analysis

For all RNA-Seq experiments, analyses were performed through a combination of KBase-(Arkin et al., 2018) and custom jupyter notebook-based methods. The data processing narrative in KBase can be found here: https://kbase.us/n/48675/70/. StringTie and DESeq2 KBase outputs are currently available in Datasets S5 and S6 (https://doi.org/10.6084/m9.figshare.12185031). Briefly, Illumina reads were trimmed using Trimmomatic v0.36 (Bolger et al., 2014) and assessed for quality using FASTQC. Trimmed reads were subsequently mapped to the *S*. Typhimurium LT2 along with PSLT genome (NCBI Accession: AE006468.2 and AE006471.2 respectively) with HISAT2 v2.1.0 (Kim et al., 2019). Alignments were quality-assessed with BAMQC. From these alignments, transcripts were assembled and abundance-estimated with StringTie v1.3.3b (Pertea et al., 2015). Tests for differential expression were performed on normalized gene counts by DESeq2 (negative binomial generalized linear model) (Love et al., 2014). Additional analyses for all experiments were performed in Python3 and visualized employing matplotlib and seaborn packages. Conservative thresholds were employed for assessing differentially expressed genes. Conclusions were considered differentially expressed if they possessed a Bonferoni-corrected p-value below a threshold of 0.001 and an absolute log2 fold change greater than 2. Assembled transcripts from StringTie and differential expression from RNA-Seq analyses can be found in Datasets S5 and S6 respectively.

### Genome sequencing of SARA collection

We sequenced the 21 reference *S*. Typhimurium genomes(Beltran et al., 1991)using standard molecular biology protocols. Briefly, we grew up all 21 strains to stationary phase in LB media. We then extracted gDNA using the DNeasy Blood and Tissue kit (Qiagen). Illumina library preparation was performed by the Functional Genomics Laboratory (FGL), a QB3-Berkeley Core Research Facility at UC Berkeley. Sequencing was performed at the Vincent Coates Sequencing Center, a QB3-Berkeley Core Research Facility at UC Berkeley on a HiSeq4000 using 100PE reads. We used Unicycler with default parameters(Wick et al., 2017) to do a reference based assembly from closely related *Salmonella* strains.

### Bioinformatic analysis of SARA collection genomes

Predicted genes in 24 *Salmonella typhimurium* genomes were classified in families of homologous genes by PPanGGoLiN(Gautreau et al., 2020). Gene clusters encoding LPS core oligosaccharide and O-specific antigen (OSA) biosynthetic enzymes were identified in the *Salmonella* genomes by search for gene families containing characterized LPS and OSA biosynthesis genes of LT2 strain (STM2079-STM2098, STM3710-STM3723)(Heinrichs et al., 1998; Seif et al., 2019). O-antigen modification genes were identified by DIAMOND similarity search(Buchfink et al., 2015) with characterized LT2 proteins OpvA (STM2209), OpvB (STM2208), GtrA (STM0559, STM4204), GtrB(STM0558, STM4205)(Broadbent et al., 2010) using blastp command with --very-sensitive option. Restriction/modification genes were identified by DIAMOND similarity search with 78008 proteins from REBASE database(Roberts et al., 2015). Point mutations in LPS and OSA biosynthesis enzymes were identified by running a command-line application for TBLASTN search(Camacho et al., 2009) of LT2 proteins vs. genome sequences of 23 *Salmonella typhimurium* genomes.

### Phylogenetic analysis

To estimate phylogenetic relationships between genomes of our collection of *Pseudomonas* spp. strains, we identified a set of 120 bacterial marker genes with GTDB-Tk toolkit(Chaumeil et al., 2019). Only 115 marker genes were found in single copy in each of the 24 genomes studied. Gene sequences of those 115 markers were aligned by MAFFT v7.310(Katoh and Standley, 2014) with *--auto* option, and the resulting 88 alignments were concatenated into a single multiple sequence alignment. A phylogenetic tree was reconstructed from the multiple alignment using the maximum likelihood method and generalized time-reversible model of nucleotide substitution implemented in the FastTree software v2.1.10(Price et al., 2010) and visualized using the Interactive Tree of Life (iTOL) online tool(Letunic and Bork, 2019).

### Prophage analysis

To determine prophage content, we submitted all contigs longer than 10kb to the PHASTER web server, culminating in 250 potential prophage regions across the 21 strains investigated(Arndt et al., 2016) (Dataset S9). These 250 identified regions were aligned against each other using nucmer(Marçais et al., 2018). Grouping and subsequent filtering of prophages was performed through network graph analysis using Gephi; prophage nodes were connected by edges representing total nucmer alignments greater than 60% alignment. Graph layout optimization was determined through layout optimization based off of equally weighted edges using the Yifan Hu algorithm. For each “cluster” of prophages and each alone prophage, a few representatives were investigated manually for prophage similarity to determine if a PHASTER-identified region (or “cluster”) was correctly identified as a prophage, yielding 84 likely prophage regions. Based on similarity to studied prophages, we assigned each “cluster” to one of “ST64B (118970_sal3-like)”, “Gifsy-1”, “Gifsy-2”, “Gifsy-3”, “Fels-1”, “P2-like”, “P22-like”, “phiKO2-like”, “SPN1S-like” classifications (Dataset S9).

Because this prophage determination was based upon a reference-based assembly(Wick et al., 2017), it was possible for some regions to mis-assemble depending on the reference genome used. So, we further validated if these prophage regions were artifacts of assembly. For each genome, we re-aligned our reads to the assembled genome using samtools and noted all regions that were not covered in BAM-alignments. We noted if prophages were either (1) split across contigs (common for “Gifsy-2”), (2) not covered by reads (noted 2 instances for P22-like prophages), (3) partially not covered by reads (common for P22-like phages, which have known mosaic sequences)(Fu et al., 2017) and (4) compared our prophage identification efforts to earlier work(Fu et al., 2017). After eliminating prophage regions that were assembly artifacts, we culminated in 74 high confidence prophage regions across the 21 SARA strains (Dataset S9).

## Acknowledgments

The authors gratefully thank Kenneth Sanderson (*Salmonella* Genetic Stock Center), Michael McClelland, Sylvain Moineau (Félix d’Hérelle Reference Center for Bacterial Viruses), Richard Calendar, Ian J. Molineux, and Jason J. Gill for sharing bacterial strains and phages and supplying valuable advice. Additionally, we would like to thank Morgan Price for helpful conversations during analysis and preparation of the manuscript.

This project was funded by the Microbiology Program of the Innovative Genomics Institute, Berkeley. The initial concepts for this project were funded by ENIGMA, a Scientific Focus Area Program at Lawrence Berkeley National Laboratory, supported by the U.S. Department of Energy, Office of Science, Office of Biological and Environmental Research under contract DE-AC02-05CH11231.

RNA sample processing and library creation was performed at Functional Genomics Lab, Vincent J. Coates Genomics Sequencing Lab, & Computational Genomics Resources Lab (University of California at Berkeley). Sequencing was performed at: Vincent J. Coates Genomics Sequencing Laboratory (University of California at Berkeley), supported by NIH S10 Instrumentation Grants S10RR029668, S10RR027303, and OD018174.

## Author Contributions

B.A.A., V.K.M., and A.P.A. conceived the project.

B.A.A. led the experimental work, analysis, and manuscript preparation.

B.A.A., V.K.M., C.Z., A.M.D. and H.L. built and characterized the RB-TnSeq library.

B.A.A. performed experiments, processed, and analyzed data.

E.B.K. provided critical reagents and advice. T.N.N., L.M.L. assembled genomes.

B.A.A., E.B.K., A.M.D., V.K.M., and A.P.A. wrote the paper.

## Data Availability

Supplementary Information can be found here: https://doi.org/10.6084/m9.figshare.12185001.v2. Complete Supplementary Datasets can be found here: https://doi.org/10.6084/m9.figshare.12185031. Supplementary Code and figure reproduction can be found here: https://doi.org/10.6084/m9.figshare.12412814.v2. All NGS reads have been deposited and made publicly accessible via the Sequence Read Archive (SRA) under Bioproject PRJNA638761: http://www.ncbi.nlm.nih.gov/bioproject/638761. Draft *S*.Typhimurium genome sequences for strains SARA1-SARA21 can be found under BioSamples SAMN17506935-SAMN17506955. Draft phage genome sequences and JGI-performed gene annotations can be found at JGI IMG under analysis projects Ga0451357, Ga0451371, Ga0451358, and Ga0451372. The RNA-Seq data processing narrative in KBase can be found here: https://kbase.us/n/48675/70/.

