## Supplementary Information 2021 for "The Genetic Basis of phage susceptibility, cross-resistance and host-range in *Salmonella*"

Benjamin A. Adler^1,2,3^, Alexey E. Kazakov^4^, Crystal Zhong^2^, Hualan Liu^4^, Elizabeth Kutter^5^, Lauren M. Lui^4^, Torben N. Nielsen^4^, Heloise Carion^2^, Adam M. Deutschbauer^4,6^, Vivek K. Mutalik^3,4*^, Adam P. Arkin^2,3,4*^

1. The UC Berkeley-UCSF Graduate Program in Bioengineering, Berkeley, California, United States
2. Department of Bioengineering, University of California, Berkeley, Berkeley, California, United States
3. Innovative Genomics Institute, University of California, Berkeley, Berkeley, United States
4. Environmental Genomics and Systems Biology Division, Lawrence Berkeley National Laboratory, Berkeley, California, United States
5. The Evergreen State College, Olympia, Washington, United States
6. Department of Plant and Microbial Biology, University of California, Berkeley, Berkeley, California, United States

* Correspondence:

Vivek K. Mutalik, Adam P. Arkin

**Text S1**

**Extended Results: RB-TnSeq LPS Mutants.**

In *Salmonella* species, LPS is well-characterized and highly diversified at O-antigen residues. Generally, LPS structure consists of 4 regions: KDO sugars covalently bonded to lipid A, the inner core, outer core, and O-antigen repeat (Figure 2A). In wild-type *S.* Typhimurium LT2 (parental strain for MS1868), the O-antigen is made up of 200 hexose monomers per polymer (Murray, Attridge, and Morona 2003). These regions vary phylogenetically, with KDO and inner core structures being conserved across many genera and outer core structures not conserved between species. O-antigen structures in particular exhibit high degrees of variability – approximately 46 O-antigen structures have been discovered in *Salmonella spp.* (Liu et al., n.d.). Within the same strain, O-antigen structures can exhibit variability; through a variety of regulatory mechanisms, O-antigen can manifest as capsular polysaccharide, vary in repeat number, and vary in sugar modifications (Gibson et al. 2006; Cota, Blanc-Potard, and Casadesús 2012; Murray, Attridge, and Morona 2003; Broadbent, Davies, and Van Der Woude 2010).

The MS1868 library included mutants in 25 genes responsible for LPS biosynthesis and transport. 15 of these genes are involved in core LPS biosynthesis (*rfaH*, *rfaC*, *rfaZ*, *yaeD*, *rfaD*, *rfaP*, *rfaE*, *rfaF*, *rfaQ*, *rfaY*, *rfaG*, *rfaB*, *galE*, *rfaI*, and *rfaK*) and 9 genes are involved in O-antigen biosynthesis (*rfaL, wzzB*, *rfbP*, *rfbK*, *rfbC*, *rfbD*, *rfbB*, *pgm*, *galE*, and *oafA*). GalE synthesizes a precursor used in both LPS core and O-antigen biosynthesis (Figure 2A). In general, a loss of function mutation in a biosynthetically upstream gene disrupts remaining LPS and O-antigen biosynthesis. (1) *rfaZ*, *rfaY*, and *rfaQ* are not required for O-antigen maturation (Klena, Ashford, and Schnaitman 1992), (2) *rfaB* mutants yield strains with a heterogenous LPS O-antigen phenotype (Wollin et al. 1983), (3) *yaeD* mutants in *E. coli* can form heptose-less LPS (Kneidinger et al. 2002), and (4) *oafA* mutants do not otherwise affect O-antigen maturation (Hauser et al. 2011). In general, LPS requirements for each phage were found to be consistent between fitness data and an established chemotype-defined LPS mutant panel in *S.* Typhimurium, representing 14 distinct LPS chemotypes (*Salmonella* Genetic Stock Center (SGSC), Figures S3-S11, S13-S15). An overview of the LPS requirements for phages Aji_GE, Chi, FelixO1, P22, Reaper_GE, Savina_GE, and SP6 can be seen in Figures S3 and S12. An overview of the LPS requirements for phages Br60, Ffm, and Shishito_GE can be seen in Figure S4. As a resource, an in-depth discussion of these results is continued below.

**Extended Results: O-Antigen Requiring Phages P22 SP6, Reaper_GE, and Savina_GE**

Four of the phages tested (P22, SP6, Reaper_GE, Savina_GE) displayed strict requirements for full O4[5] antigen biosynthesis despite being unrelated (Figures S3BC and S12, Table 1). This was expected for both P22 and SP6 since the O4[5] antigen is the established receptor for these phages (Wright, McConnell, and Kanegasaki 1980; Steinbacher et al. 1997; Tu et al. 2017). SP6 and P22 had the most stringent LPS and O-antigen requirements, requiring 19 and 18 of the 25 LPS and O-antigen biosynthesis genes respectively across liquid and solid fitness experiments. Relative to P22, SP6 additionally required *yaeD* activity, suggesting that it is less infective on heptose-less LPS (Kneidinger et al. 2002). Although *rfaZ* and *rfaQ* gave high mean fitness scores for both phages, these scores were only high when the expected insertion was oriented against transcription (example shown in Figure S1C), indicating that these fitness values were likely due to polar, transcription-disrupting effects on downstream genes. These results were largely consistent with a related genome-wide screen against P22 infection (Bohm et al. 2018). Compared to this study, host-factor requirement differences for P22 were largely attributed to differences in library coverage. Because the coverage of each library is different, each library can interrogate unique genes that the other can not. Due to better coverage in our library, our screens identified *rfaC*, *rfaD*, and *rfaE*. Due to lower coverage in our library, we were unable to analyze effects of mutants for several genes in O-antigen biosynthesis (highlighted in Figure 2A). Despite being analyzed in both studies, our study additionally identified *rfaP* as a novel host-factor for P22 (as well as SP6).

Unlike P22 and SP6, Reaper_GE and Savina_GE are novel phages investigated in this study and had no prior known host-requirements. The only genes Reaper_GE and Savina_GE required in this study were 16 and 15 of the 25 LPS and O-antigen biosynthesis genes respectively across liquid (both) and solid (Reaper_GE only) fitness experiments. Some additional host-requirement differences were noted between Reaper_GE and Savina_GE versus P22 and SP6. For instance, strains with *yaeD*, *pgm*, and *rfbB* (responsible for inner-core heptose biosynthesis and O-antigen precursor biosynthesis (Figure 2A)) disruptions were sensitive to Reaper_GE or Savina_GE phages. Additionally, the only *rfaP* disruptions that were fit against Reaper_GE and Savina_GE were oriented against transcription and are likely polar effects. This absolute requirement for intact O-antigen including a complete LPS core was validated for Reaper_GE in our LPS chemotype panel (Figure S7). Based on these results, we propose O-antigen as the receptor for Reaper_GE phage.

Unlike Reaper_GE, Savina_GE did not require *rfaI* (responsible for the second glucose addition to the outer core (Figure 2A)) and *rfaH* (responsible for activation of *rfa* and *rfb* operons). We found that fitness scores for genes changed in magnitude by position in the proteins’ role in LPS biosynthesis. For instance, we found that O-antigen mutants displayed the strongest fitness in the presence of Savina_GE, followed by inner core mutants, and then outer core mutants (Figure S3B, Dataset S4), suggesting differential host interaction with LPS and O-antigen moieties. Unlike all other phages investigated here, solid assays yielded little selection pressure against Savina_GE, suggesting generally poor infection of Savina_GE to wild-type MS1868. We further investigated this fitness pattern on our LPS chemotype panel and found results consistent with our BarSeq data; phage Savina_GE was most infective against strains with an incomplete outer core, but less so against strains without O-antigen or strains missing outer core entirely (Figure S9). Based on this data, it appears that phage Savina_GE employs an infection strategy similar to phage PVP-SE1(Santos et al. 2011). Potentially, Savina_GE preferentially employs LPS as a receptor, but branched LPS residues such as those added by *rfaK* and O-antigen biosynthesis hinder adsorption.

**Extended Results: FelixO1 LPS Requirements**

FelixO1 gave results consistent with literature identifying outer core GlcNAc, the biosynthetic product of RfaK, as FelixO1’s primary receptor (Lindberg and Hellerqvist 1971). In addition, mutants in 12 of the LPS biosynthesis genes conferred resistance against FelixO1 (*rfaH*, *rfaC*, *yaeD*, *rfaD*, *rfaP*, *rfaE*, *rfaF*, *rfaG*, *rfaB*, *galE*, *rfaI*, and *rfaK*) in both liquid and solid fitness experiments. High fitness score of *yaeD* (responsible for inner-core heptose biosynthesis), suggested the importance of inner core integrity for FelixO1 infection (Kneidinger et al. 2002). We also found an additional 61 non-LPS genes important in FelixO1 infection, which will be discussed below (see “Discovery of Novel Cross-Resistant Genotypes Between Diverse Phages” in the main text).

**Extended Results: LPS Requirements of Protein Receptor Phages S16, Chi, and Aji_GE**

S16, Chi, and Aji_GE all likely employ protein-based receptors (see Extended Results: Protein Receptor Requirements of Protein Receptor Phages S16, Chi, and Aji_GE), but also each had some degree of reliance on LPS. *rfaI* and *rfaG* mutants provided small, but significant, fitness benefits against phage S16 (Figure S3). Plaque assays against defined LPS chemotypes suggested decreased infectivity due to *rfaF* and *rfaG* mutants (Figure S10), but overall, these results are consistent with literature suggesting that inner core *Salmonella* LPS assist in, but are not strictly required for, S16 adsorption (Marti et al. 2013). Mutants in *rfaH*, *yaeD*, *rfaP*, *rfaQ*, *rfaY*, *rfaG*, and *pgm* all provided fitness benefits against phage Chi, results not before observed for Chi. Nonetheless, it is likely that these mutants primarily impact motility, thus reducing Chi infectivity (see Extended Results: Protein Receptor Requirements of Protein Receptor Phages S16, Chi, and Aji_GE) (Toguchi et al. 2000); additionally, homologs of *rfaH*, *rfaP*, *rfaQ*, *rfaY*, and *rfaG* are among the least motile mutants in experiments for *E. coli* BW25113 (Price et al. 2018). Since LPS disruptions induce stress responses, this pleiotropy may be stress-induced downregulation of outer membrane proteins and flagellar activity (Mutalik et al. 2020).

**Extended Results: Protein Receptor Requirements of Protein Receptor Phages S16, Chi, and Aji_GE**

Phages S16, Aji_GE, and Chi showed dependence on outer protein structures in addition to LPS. Prior genetic and biochemical characterization of T4-like phage S16 identified OmpC as the primary receptor (Marti et al. 2013). Concordantly, in our screens, *ompC* loss of function mutants were highly fit against S16. In addition, mutants of positive regulators of *ompC* expression, *ompR* and *envZ* (Mizuno et al. 1988), were enriched against S16 (Figure S16AB). Our screens are consistent with earlier findings of OmpC as the primary receptor with influence from LPS for S16, similar to similar screens against related phage T4 (Figures 1B, S3BC, and S16ABC) (Marti et al. 2013; Mutalik et al. 2020). Because loss-of-function of OmpC yields S16 phage resistance and OmpC has been implicated in antibiotic resistance in *Salmonella* and related species (Pagès 2004; Fernández and Hancock 2012; Ghai and Ghai 2018), there may be synergetic effects through use of S16 as an antibiotic adjuvant.

For T5-like phage Aji_GE, we find *fepA* (that encodes a TonB-dependent enterobactin receptor) as a high-scoring host-factor in addition to *oafA*. As FepA is known to form a complex with TonB and function as a receptor for some T5-like phages, such as phage H8 (Rabsch et al. 2007), we believe the FepA-TonB complex is the primary receptor for phage Aji_GE (Figure S16AC). As our RB-TnSeq library lacked *tonB* mutants, we created *fepA* and *tonB* deletions to validate the role of FepA and to assess the requirement of TonB. Indeed, individual *fepA* or *tonB* deletion mutants gave resistance to Aji_GE, confirming their essentiality for Aji_GE infection (Figures S16-S17). These results indicate that, potentially similar to phages T5 and SPC35, phage Aji_GE employs LPS modifications (here, added by OafA) to enhance infection (Heller and Braun 1982; Kim and Ryu 2012) and gain access to the FepA-TonB complex for efficient infection.

Bacteriophage Chi is a model flagellar-binding bacteriophage that employs flagellar activity to approach the *S.* Typhimurium outer membrane (Schade, Adler, and Ris 1967). Because flagellar assembly and activity is dependent on the concerted activity of around 50 genes (for a review on enteric flagella, see Macnab et al. (2003) (Macnab 2003)), we postulated that a systems-level view may provide efficient insight into the host-requirements for Chi-like phage versus individual genetic mutant studies. As expected, 36 genes across *flg*, *flh*, *fli*, and *mot* operons responsible for flagellar biosynthesis and regulation gave strong fitness scores, implicating the importance of flagellar biosynthesis for phage Chi infection (Figure S16A).

In addition, we found a number of flagellar activity-modulating factors important against phage Chi infection. For example, we observed *cheZ* mutants fit against Chi phage (Figure S18). CheZ functions by dephosphorylating CheY-phosphate, biasing flagella counterclockwise towards smooth swimming (Macnab 2003). Putatively, loss of CheZ activity biases flagellar activity clockwise, leading to tumbling and subsequent Chi resistance, consistent with CheY activity studies and Chi phage (Samuel et al. 1999). In addition, several mutants in guanosine penta/tetraphosphate ((p)ppGpp) biosynthesis and metabolism were enriched in Chi infection experiments, potentially due to impacts on motility (Christen et al. 2016). Other top-scoring mutants against Chi infection include *nusA*, *tolA*, and *cyaA,* that were earlier found to greatly decrease *E. coli* motility (Price et al. 2018), suggesting that indirect motility defects provide numerous routes to resist flagella-dependent phages. Because phase I flagellin, *fliC*, show high fitness scores, but phase II flagellin, *fljB*, and *fliC*’s repressor, *fljA*, did not, Chi infection could be specific to phase I flagellar phenotypes (Ikeda et al. 2001). However, further studies are needed to dive deeper and mechanistically confirm this result.

As high-throughput genetic screens to infer interactions between bacteriophage and their hosts grow more commonplace, we expect to see further enrichments in our understanding of bacteriophage adsorption requirements. For instance, we doubt that dual LPS + protein receptor combinations are unique to S16, Aji_GE, and Chi - many such combinations likely remain to be discovered as more phage are characterized. Additionally, many Chi-like bacteriophage variants exist, some of which depend on different flagellar rotations or other flagellin filaments (Christen et al. 2016; Choi et al. 2013). Due to the pooled nature of RB-TnSeq experiments, we are able to characterize in a single experiment what would have otherwise taken over 75 individual mutant experiments for some of these phages.

**Extended Results: Uncovering Host-Factors of Rough-LPS Requiring Phages Br60, Ffm, and Shishito_GE**

Br60, Ffm, and the newly isolated Shishito_GE phages are capable of infecting *Salmonella* with rough-phenotype-LPS (this phenotype hereon referred to as “rough-LPS”), but not smooth phenotype LPS (ie wild-type for MS1868) (Figures S13-S15). Against O-antigen positive strains, such as our library base strain, MS1868, none of these phages successfully infect as they are occluded from their native receptor by the O-antigen structure. However, they can infect strains without O-antigen (Wilkinson, Gemski, and Stocker 1972) (Figures S4, S13-S15). Thus, the resistance landscape between O-antigen requiring phages (for instance P22, Reaper_GE, or SP6) and rough-LPS requiring phages (Br60, Ffm, and Shishito_GE) presents an evolutionary trade-off through collateral sensitivity. In other words, many mutants resistant to O-antigen requiring phages display increased sensitivity to the rough-LPS requiring phages.

As expected with our MS1868 library primarily consisting of O-antigen positive mutants, the vast majority of gene disruptions in MS1868 showed no significant fitness benefit against these phages. However, we noticed strong fitness defects in many of the LPS and O-antigen mutants in our library (Dataset S4), consistent with optimal adsorption and infection in O-antigen-defective *Salmonellae*.

Many lab model strains of *E. coli* lost their ability to produce O-antigen, but have a similar inner and outer core LPS structure to *S.* Typhimurium (Heinrichs, Yethon, and Whitfield 1998). Thus we postulated and confirmed that these phages may additionally infect strains such as *E. coli* K-12 BW25113 (BW25113) (Figure S4C). Thus, we repeated experiments for these 3 phages using an earlier reported *E. coli* RB-TnSeq library as a model rough-LPS library analog to the MS1868 library (Wetmore et al. 2015; Mutalik et al. 2020). Consistent with related phages T3 and T7 from earlier library analyses, few host-factors were required for infection. All 3 phages showed slightly different LPS requirements for infection. Against Ffm and Shishito_GE, mutants in *gmhA*, *hldE*, *hldD*, and *waaC* were phage-resistant (*ghmA*, *rfaE*, *rfaD*, *rfaC* MS1868 homologs respectively). Ffm additionally required *waaG* (*rfaG* MS1868 homolog) for infectivity. However, no LPS or protein mutants were fit against Br60. *trxA* mutants were fit against all three rough-LPS requiring phages (Figure S4D, Dataset).

To confirm that these results were extensible to *S.* Typhimurium, we assayed infectivity against our LPS mutant collection (SGSC, Figures S13-S15). Indeed, the results observed against the *S.* Typhimurium LPS panel were perfectly analogous to the fitness experiments observed against the BW25113 library, with the exception of *rfaB*, which has a known heterogenous, smooth phenotype in *S.* Typhimurium and likely occludes adsorption (Wollin et al. 1983). *S.* Typhimurium homologs of host-factors inferred from the BW25113 library experiments were important for infection of these phages. Ffm required function of *rfaC*, *rfaD*, *rfaG*, *rfaB*, and loss of O-antigen (Figure S13). Shishito_GE required function of *rfaC*, *rfaD*, *rfaB*, and loss of O-antigen (Figure S14). Br60 only required loss of O-antigen (Figure S15). From our *E. coli* and *S.* Typhimurium LOF libraries and *S.* Typhimurium LPS mutant collection results, we conclude that these related T7-like bacteriophages require distinct LPS moieties, providing additional resolution to the putative receptors of these phages. Br60 emerged from these experiments as a particularly interesting phage due to the inability to detect important host-factors other than *trxA* from the saturated BW25113 transposition library; potentially this phage binds to LPS KDO sugars. These LPS residues are synthesized by enzymes encoded by essential genes, making host-side loss-of-function a futile way for a pathogen to escape this phage. Because the loss of function cross-resistance with O-antigen requiring phages (Figure S3BC) is minimized with Br60, Br60 appears to be a better candidate for exploiting collateral sensitivity than Ffm or Shishito_GE if applied alongside an O-antigen requiring phage.

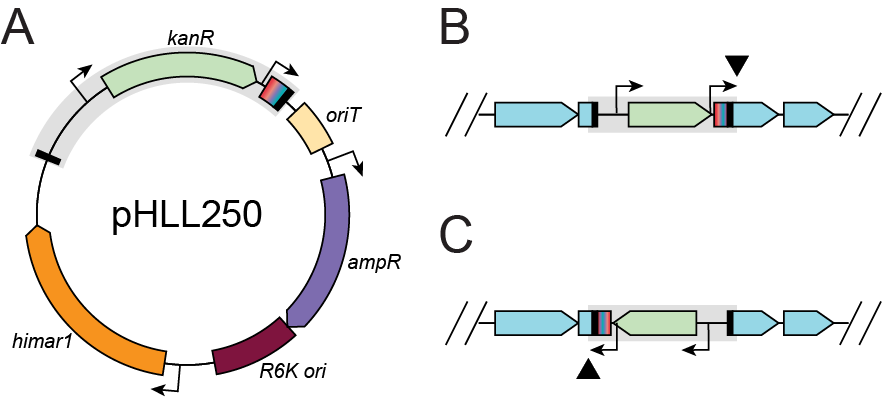

**Figure S1. Overview of RB-TnSeq Vector pHLL250.** A. Vector diagram of transposition donor vector, pHLL250. The transposable region is highlighted in gray, containing upstream and downstream inverted repeat sites (black), kanamycin resistant marker, kanR (green), and a library of N20 DNA barcodes with flanking BarSeq priming regions (rainbow). In front of the barcode is a T7 promoter to minimize polar effects in the forward orientation (see B). The vector also contains ampicillin resistance marker, ampR (purple), host-limited origin of replication R6K (maroon), conjugative transfer element, oriT (peach), and transposon himar1 (orange). For more details on how RB-TnSeq and how the parts shown here work, please refer to (20, 21). The exact sequence of this vector can be found at https://benchling.com/s/seq-gtuLW5A04BN23wfpxFGo. B. Example of forward orientation insertion in the Salmonella genome (genes shown in turquoise). The transposed element is highlighted in gray and the recorded position from RB-TnSeq corresponds to the black triangle. Per insertion event, the rainbow barcode is represented by a single variant. C. Example of reverse orientation insertion in the Salmonella genome (genes shown in turquoise). The transposed element is highlighted in gray and the recorded position from RB-TnSeq corresponds to the black triangle. Per insertion event, the rainbow barcode is represented by a single variant.

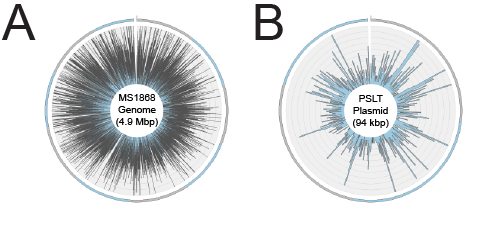

**Figure S2. Overview of RB-TnSeq Insertions in S. Typhimurium MS1868.**

Insertion density maps of new RB-TnSeq library in *S.* Typhimurium MS1868 mapped against *S.* Typhimurium LT2 reference genome (A) and PSLT plasmid (B). The gap in insertion density in quadrant III against the *S.* Typhimurium LT2 reference genome is attributed to the absence of prophage Fels2 in MS1868 relative to the LT2 reference genome(Graña, Youderian, and Susskind 1985). Input data for Figure S2 can be found in Supplementary Code - Supplemental_Figure_S2A.

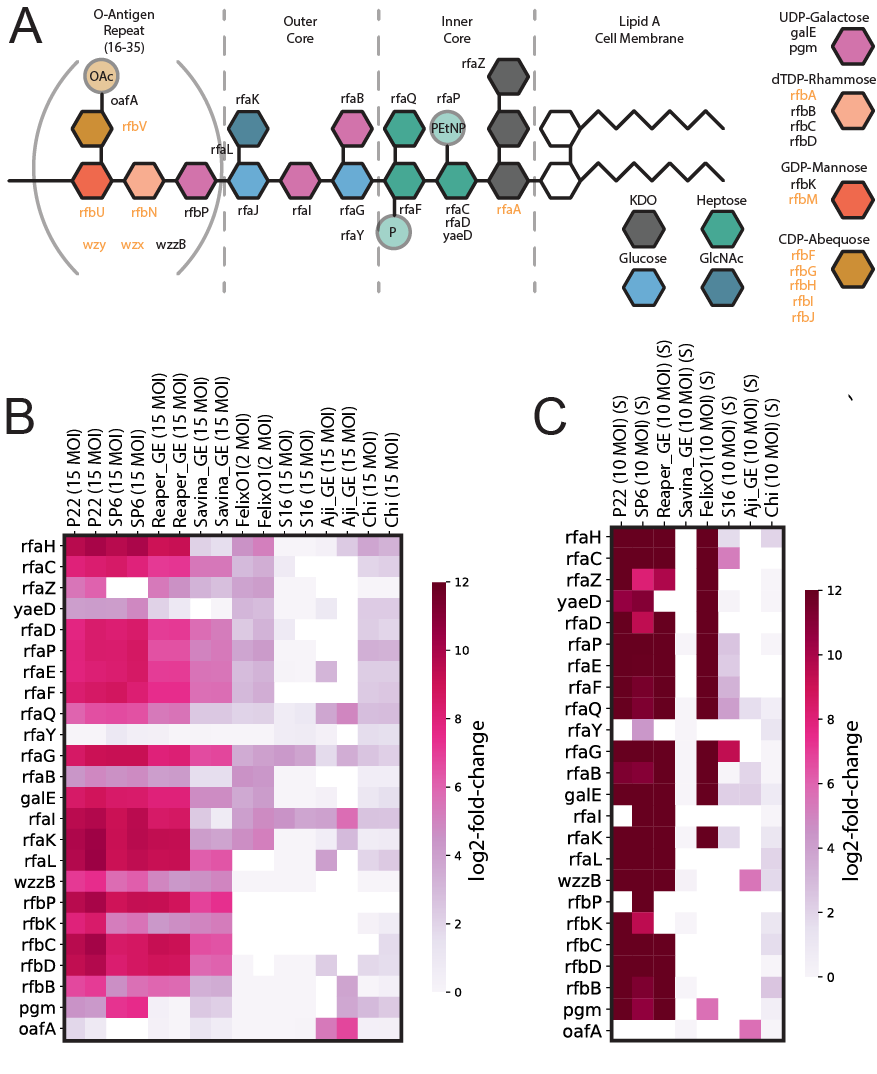

**Figure S3. Diverse LPS-Specificity Requirements for Bacteriophages Characterized in this Study.**

(A) Overview of O5 *S.* Typhimurium LPS and O-antigen biosynthesis as characterized previously. The four sugars in brackets comprise the O-antigen, which repeats 16-35 times per LPS molecule under standard growth conditions. Key for non-essential LPS and O-antigen precursor biosynthesis genes are described to the right. Genes covered in our library and used for analysis are written in black. Genes not covered in our library, and thus not analyzed in this study are written in orange. (B) Heatmap overview of gene fitness data for LPS-biosynthesis genes covered in the MS1868 RB-TnSeq library under liquid growth conditions. Genes with under 25 BarSeq reads in the phage samples had their fitness values set to 0 for visualization purposes. **(**C) Heatmap overview of gene fitness data for LPS-biosynthesis genes covered in the MS1868 RB-TnSeq library under solid media fitness conditions. Genes with under 25 BarSeq reads in the phage samples had their fitness values set to 0 for visualization clarity. Input data for Figures S3B and S3C are found in Dataset S4 and can be recreated using Supplementary Code - Supplemental_Figure_S3BC.

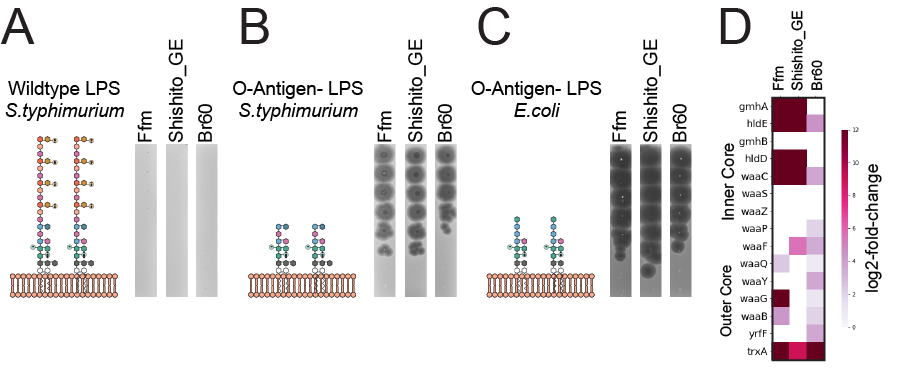

**Figure S4. Characterization of Ffm, Shishito_GE, and Br60 Adsorption Through Comparative RB-TnSeq Library Analysis.**

(A) Overview of LPS structure in smooth, O5 *S.* Typhimurium (key in Figure S3A) and representative plaque assays of Ffm, Shishito_GE, and Br60. Presumably rough-LPS requiring phage cannot access their true receptor due to presence of O-Antigen. (B) Overview of LPS structure in rough LPS, O5 *S.* Typhimurium (key in Fig S3A) (*rfaL* mutant). Rough-LPS *S.* Typhimurium strains (lacking O-antigen) show efficient phage infection. (C) Overview of rough-LPS structure in, *E. coli* BW25113 (same key in Fig S3A), which shares most of the residues in inner and outer core as rough-LPS *Salmonellae*. Despite host phylogenetic distance, we observe that Ffm, Shishito_GE, and Br60 phages efficiently infect *E. coli* BW25113. (D) Heatmap overview of functional data for LPS-biosynthesis genes covered in the *E. coli* BW25113 RB-TnSeq library under short-time-adsorption conditions on solid media for rough-LPS requiring phages Ffm, Shishito_GE, and Br60. Genes with under 25 BarSeq reads in the phage samples had their fitness values set to 0 for visualization purposes. Input data for Figure S4D are found in Dataset S4 and can be recreated using Supplementary Code - Supplemental_Figure_S4D.

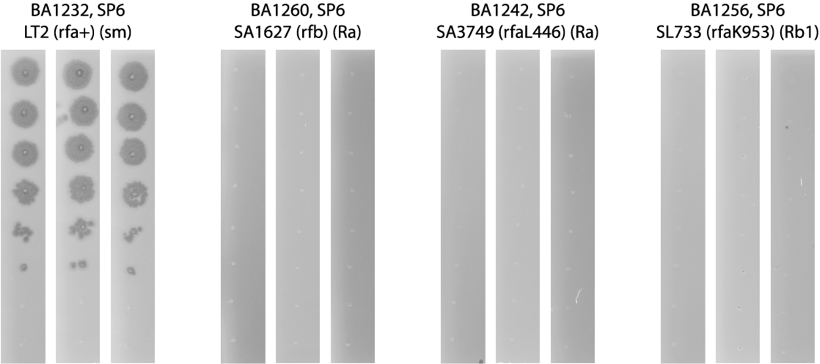

**Figure S5. LPS Chemotype Validations for Phage SP6**

Plaque assays for phage SP6 against a subset of *Salmonella* LPS mutants. Chemotypes for these strains have been identified previously and were supplied by the *Salmonella* Genetic Stock Center. Three biological replicates from unique days are shown.

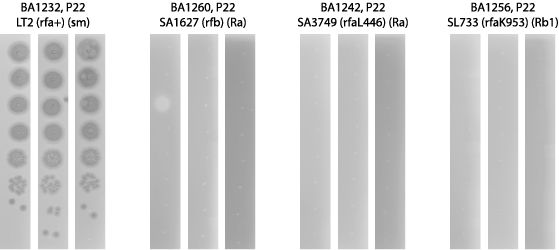

**Figure S6. LPS Chemotype Validations for Phage P22.**

Plaque assays for phage P22 against a subset of *Salmonella* LPS mutants. Chemotypes for these strains have been identified previously and were supplied by the *Salmonella* Genetic Stock Center. Three biological replicates from unique days are shown.

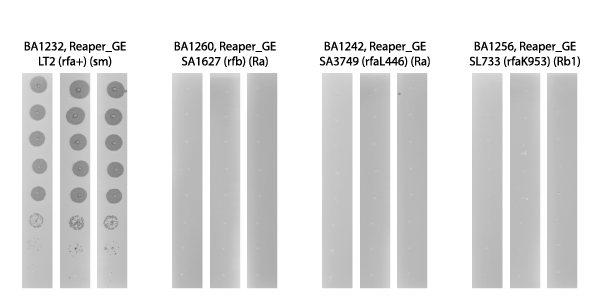

**Figure S7. LPS Chemotype Validations for Phage Reaper_GE_8C2.**

Plaque assays for phage Reaper_GE_8C2 against a subset of *Salmonella* LPS mutants. Chemotypes for these strains have been identified previously and were supplied by the *Salmonella* Genetic Stock Center. Three biological replicates from unique days are shown.

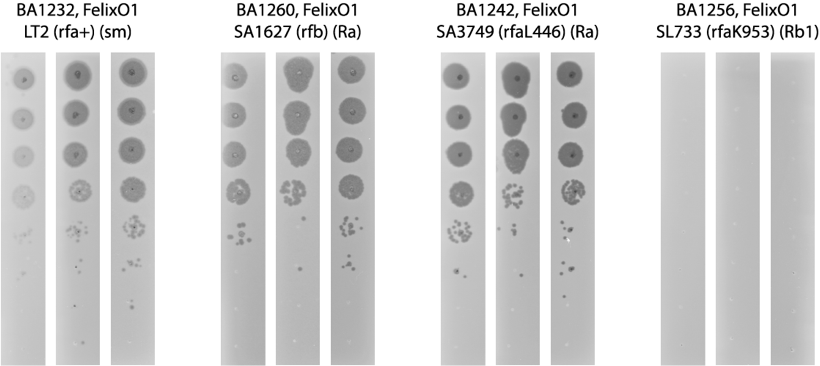

**Figure S8. LPS Chemotype Validations for Phage FelixO1.**

Plaque assays for phage FelixO1 against a subset of *Salmonella* LPS mutants. Chemotypes for these strains have been identified previously and were supplied by the *Salmonella* Genetic Stock Center. Three biological replicates from unique days are shown.

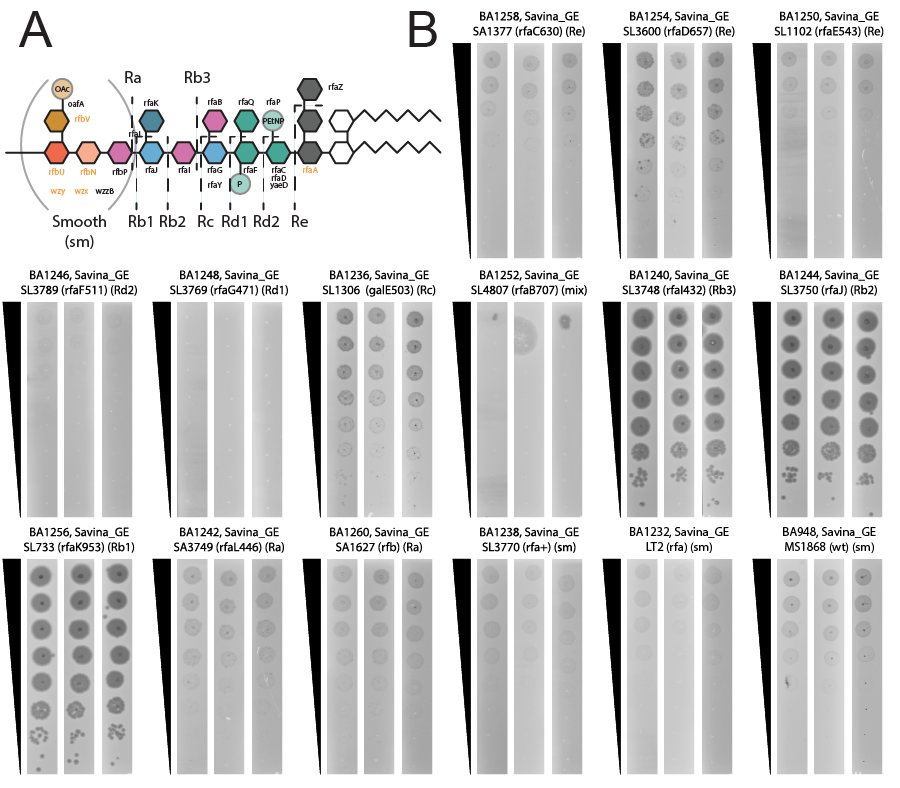

**Figure S9. LPS Chemotype Specificity Tests for Phage Savina_GE_6H2.**

A. Overview of LPS structure in *S. typhimurium* LT2 with chemotype annotations depicted.

B. Plaque assays for phage Savina_GE_6H2 against a panel of *Salmonella* LPS mutants. Chemotypes for these strains have been identified previously and were supplied by the *Salmonella* Genetic Stock Center. Three biological replicates from unique days are shown.

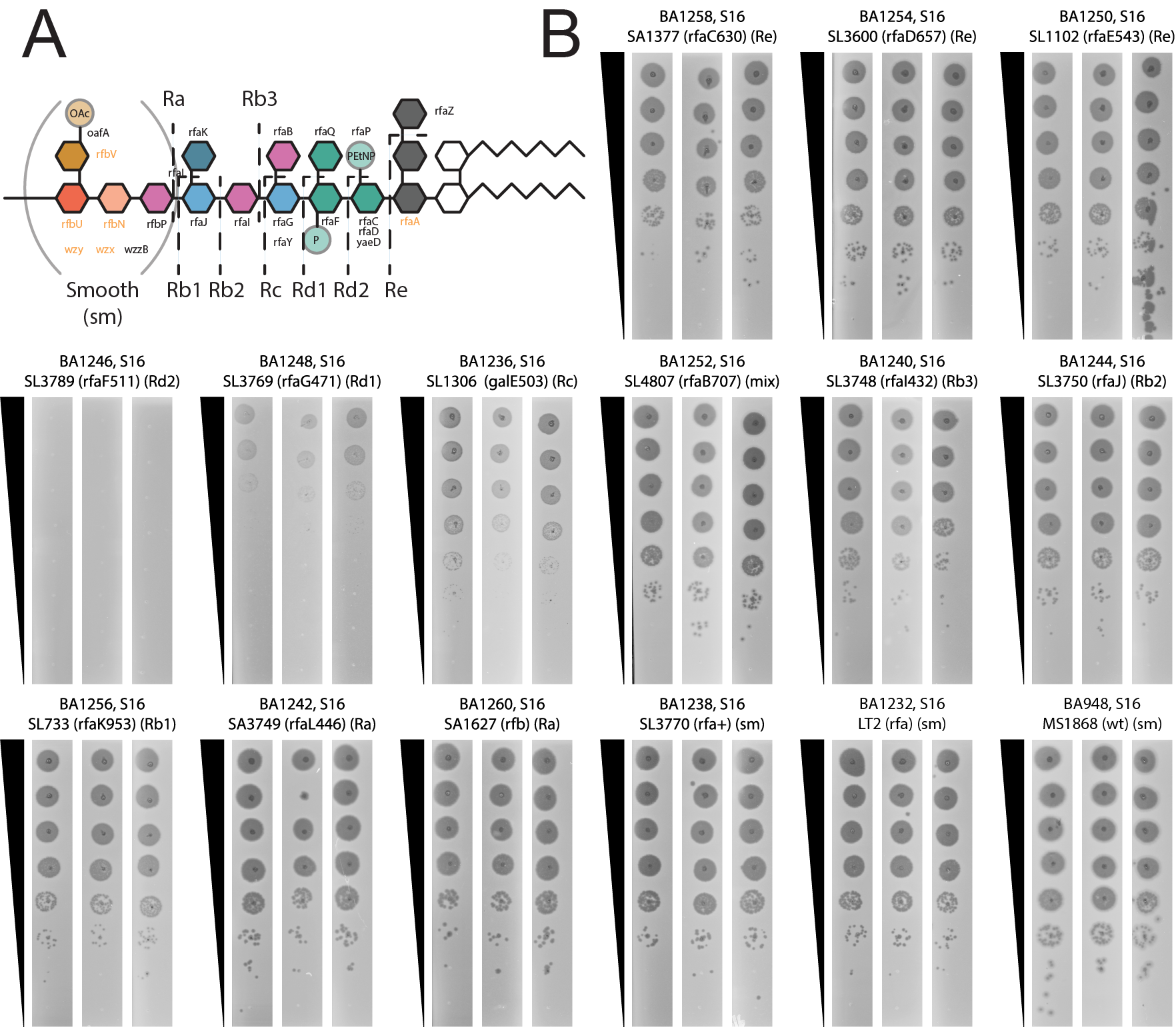

**Figure S10. LPS Chemotype Specificity Tests for Phage S16.**

A. Overview of LPS structure in *S.* Typhimurium LT2 with chemotype annotations depicted.

B. Plaque assays for phage S16 against a panel of *Salmonella* LPS mutants. Chemotypes for these strains have been identified previously and were supplied by the *Salmonella* Genetic Stock Center. Three biological replicates from unique days are shown.

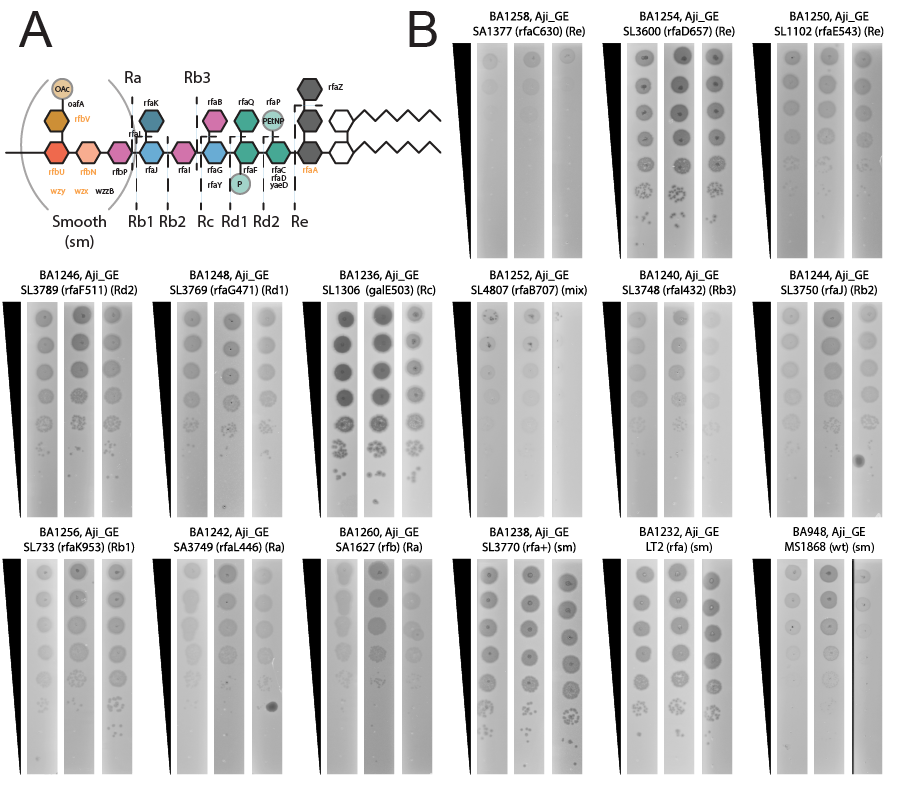

**Figure S11. LPS Chemotype Specificity Tests for Phage Aji_GE_EIP16.**

A. Overview of LPS structure in *S.* Typhimurium LT2 with chemotype annotations depicted.

B. Plaque assays for phage Aji_GE_EIP16 against a panel of *Salmonella* LPS mutants. Chemotypes for these strains have been identified previously and were supplied by the *Salmonella* Genetic Stock Center. Three biological replicates from unique days are shown.

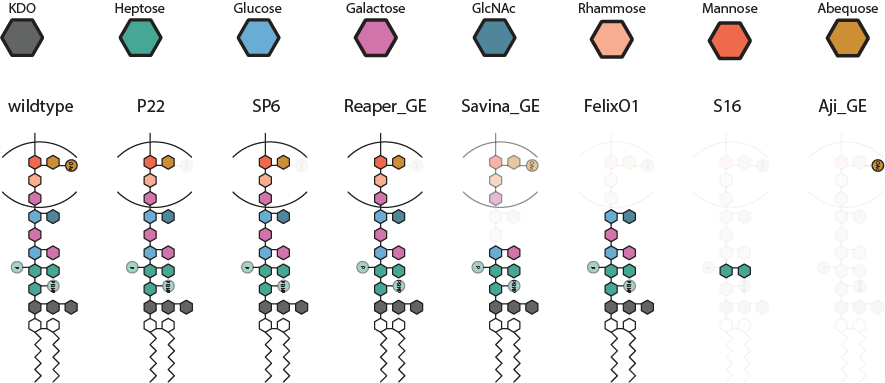

**Figure S12. Diversity of LPS Phage Resistance Factors**. Rendition of LPS-receptor for different *Salmonella* bacteriophages based on RB-TnSeq fitness values (Fig 2BC) and validations from a defined chemotype panel (S2-S8 Figs). For the modification catalyzed by OafA, a new gene replacement mutant was employed. Opaque sugar residues are strictly required for phage infection, lighter sugar and PTM residues are not required. Color key for LPS and O-antigen precursor biosynthesis sugars are shown at the top.

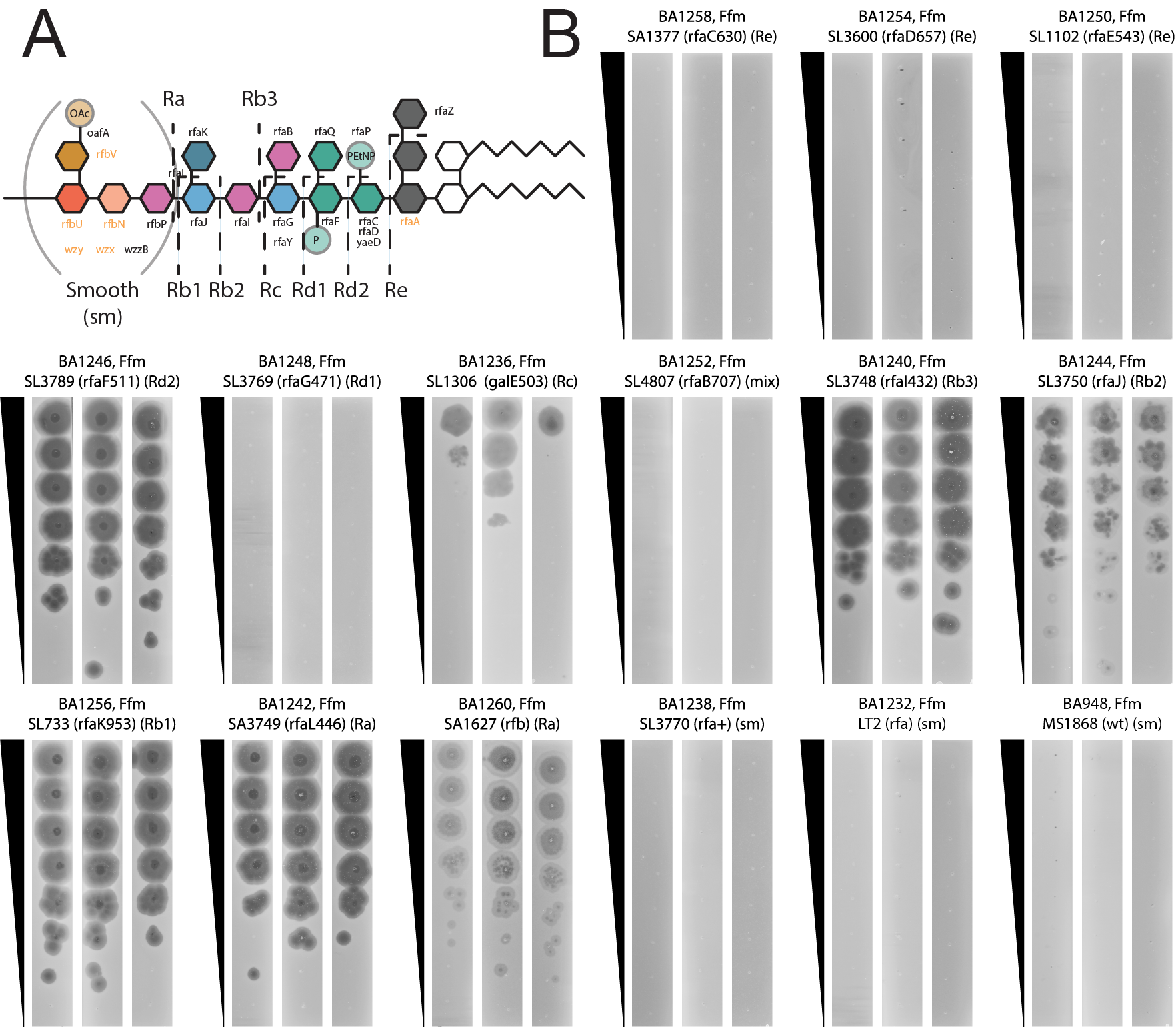

**Figure S13. LPS Chemotype Specificity Tests for Phage Ffm.**

A. Overview of LPS structure in *S.* Typhimurium LT2 with chemotype annotations depicted.

B. Plaque assays for phage Ffm against a panel of *Salmonella* LPS mutants. Chemotypes for these strains have been identified previously and were supplied by the *Salmonella* Genetic Stock Center. Three biological replicates from unique days are shown.

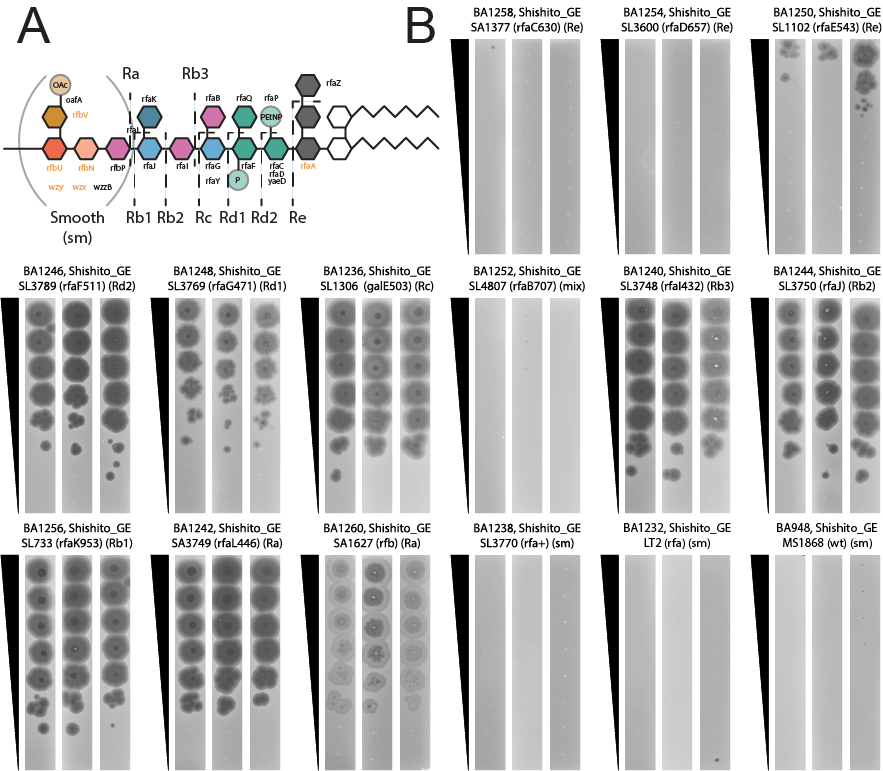

**Figure S14. LPS Chemotype Specificity Tests for Phage Shishito_GE_6F2.**

A. Overview of LPS structure in *S.* Typhimurium LT2 with chemotype annotations depicted.

B. Plaque assays for phage Shishito_GE_6F2 against a panel of *Salmonella* LPS mutants. Chemotypes for these strains have been identified previously and were supplied by the *Salmonella* Genetic Stock Center. Three biological replicates from unique days are shown.

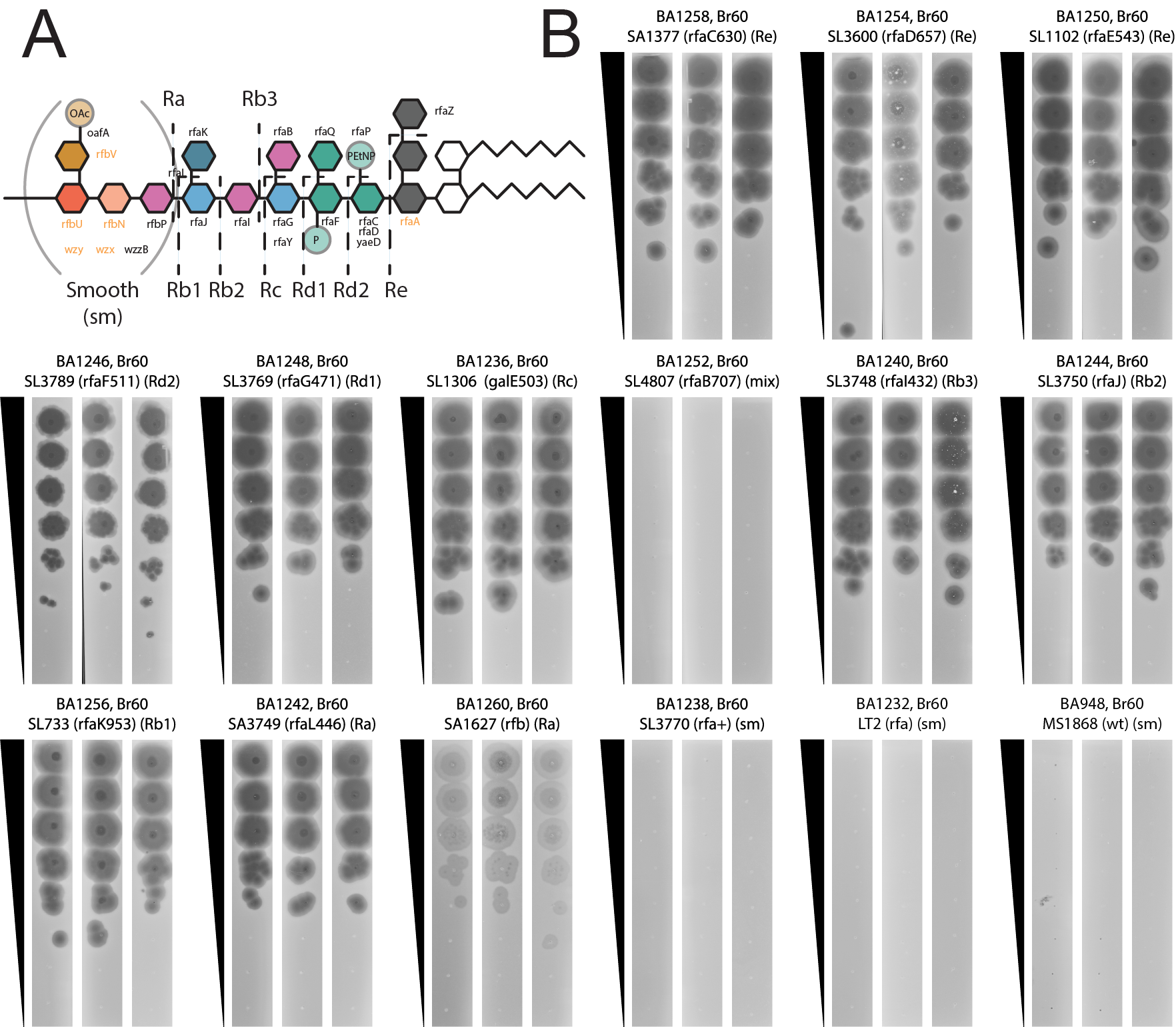

**Figure S15. LPS Chemotype Specificity Tests for Phage Br60.**

A. Overview of LPS structure in *S.* Typhimurium LT2 with chemotype annotations depicted.

B. Plaque assays for phage Br60 against a panel of *Salmonella* LPS mutants. Chemotypes for these strains have been identified previously and were supplied by the *Salmonella* Genetic Stock Center. Three biological replicates from unique days are shown.

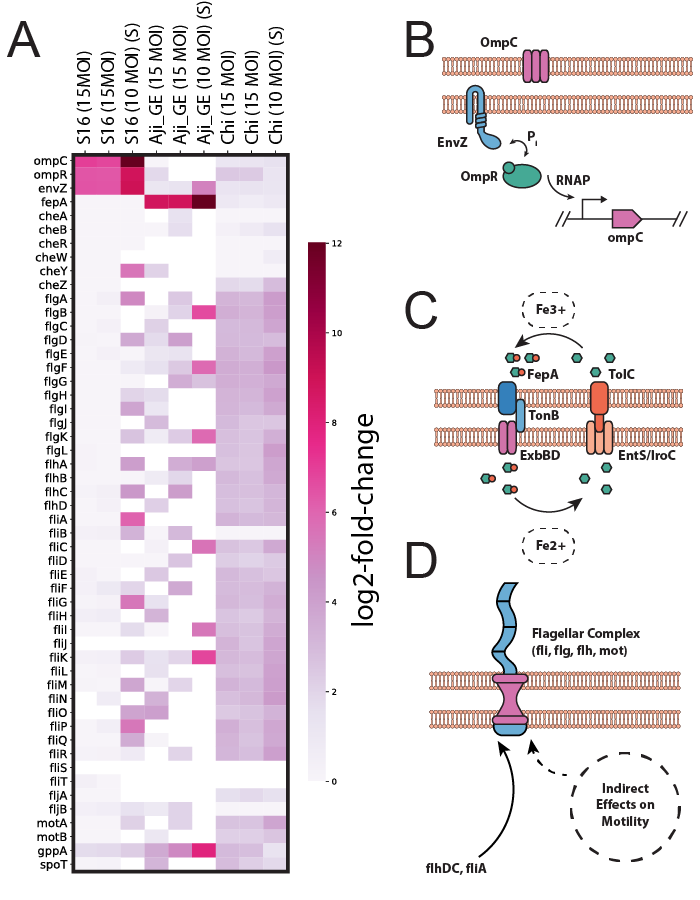

**Figure S16. Diverse protein receptors and their regulation observed for bacteriophages characterized in this study.**

(A) Heatmap overview of gene fitness data for putative protein receptors and their regulators in experiments against the MS1868 RB-TnSeq library under liquid and solid growth conditions. Noncompetitive, solid agar growth experiments are marked with a (S). Genes with under 25 BarSeq reads in the phage samples had their fitness values set to 0 for visualization purposes. (B) Schematic overview for OmpC regulation observed against phage S16. Two component system EnvZ-OmpR positively regulates expression of OmpC. If EnvZ or OmpR are disrupted, lower levels of OmpC are expressed. (C) Schematic overview for FepA, a critical host-requirement for phage Aji_GE_EIP16. FepA mediates iron scavenging through import of Fe-enterobactin and is indirectly regulated by internal iron levels. (D) Schematic overview for flagellar regulation and activity observed against phage Chi. The type I flagellar complex is assembled from proteins expressed from the multiple *flg*, *fli*, *flh*, and mot operons. Disruption of positive flagellar biosynthesis regulators *flhD*, *flhC*, and *fliA* hinders Chi infection, while disruption of negative flagellar biosynthesis regulators, *fliT* and *flgM* do not. Phase II flagellar genes *fljAB* do not appear to important for Chi infection. Other motility promoting genes such as *cheZ* are important for Chi infection as well. Input data for Figure S16A is found in Dataset S4 and can be recreated using Supplementary Code - Supplemental_Figure_S16A.

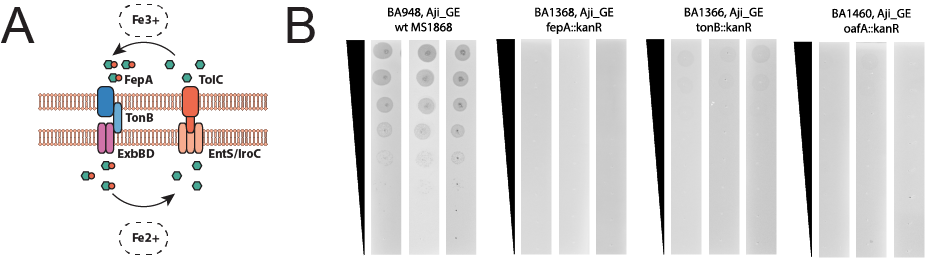

**Figure S17. Aji_GE_EIP16 Followups**

A. Overview of FepA-TonB complex in *S.* Typhimurium LT2.

B. Plaque assays for phage Aji_GE_EIP16 against putative-receptor *Salmonella* mutants. Genotypes for these strains are described. Three biological replicates from unique days are shown.

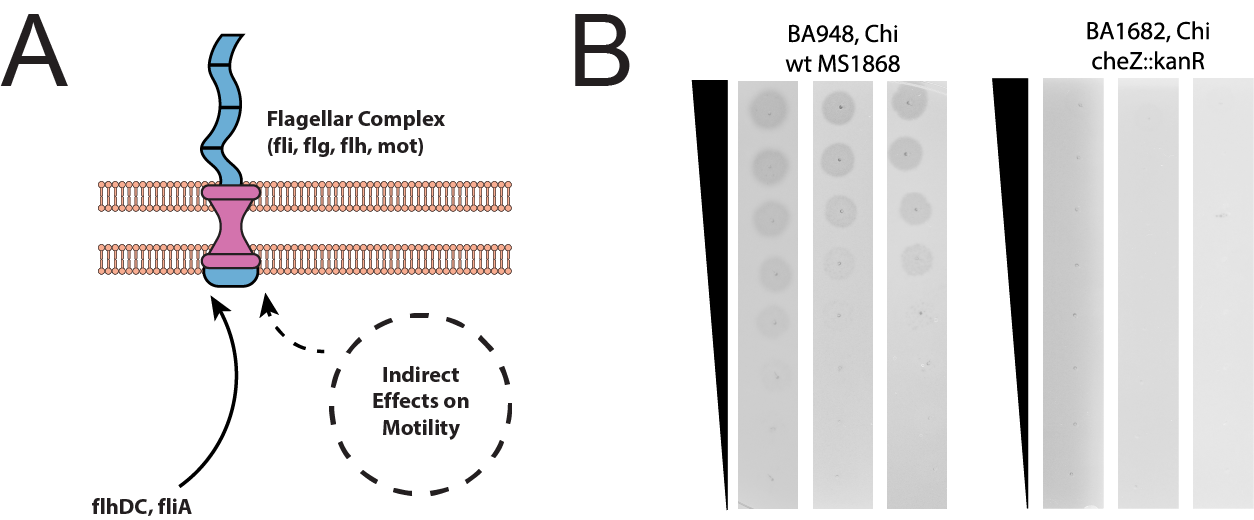

**Figure S18. Chi Followups**

A. Overview of Flagellar complex in *S.* Typhimurium LT2.

B. Plaque assays for phage Chi against wildtype and *Salmonella* *cheZ* mutant. Genotypes for these strains are described. Three biological replicates from unique days are shown.

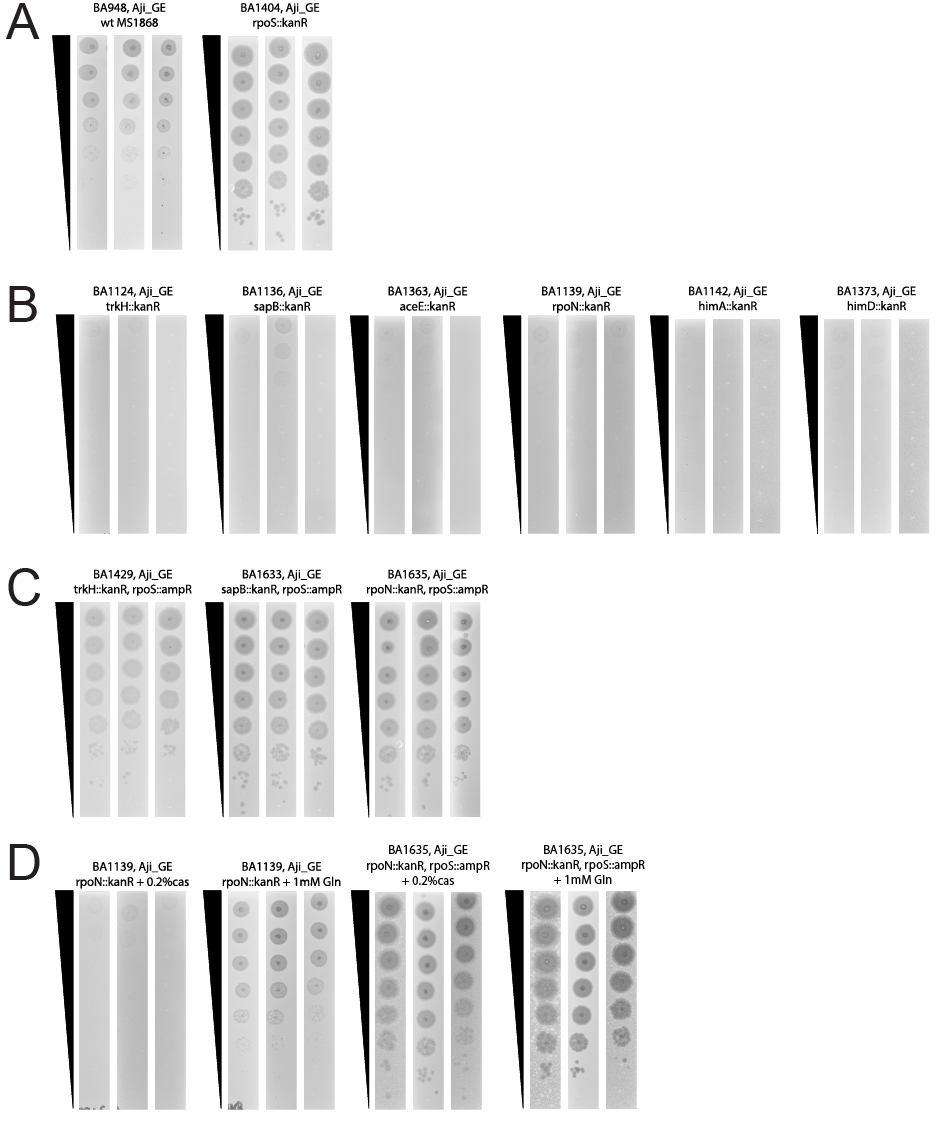

**Figure S19. Aji_GE_EIP16 Followups for Cross-Resistance Mutants.**

Plaque assays for phage Aji_GE_EIP16 against mutants identified in phage cross-resistance. In all panels, genotypes for the strains are described. Three biological replicates from unique days are shown.

A. Aji_GE_EIP16 against wildtype and *Salmonella* *rpoS* mutant.

B. Aji_GE_EIP16 plated against single deletions of cross-resistant genes.

C. Aji_GE_EIP16 plated against double deletions of cross-resistant genes and *rpoS*.

D. Aji_GE_EIP16 plated against *rpoN* and *rpoN, rpoS* mutants supplemented with casamino acids or glutamine.

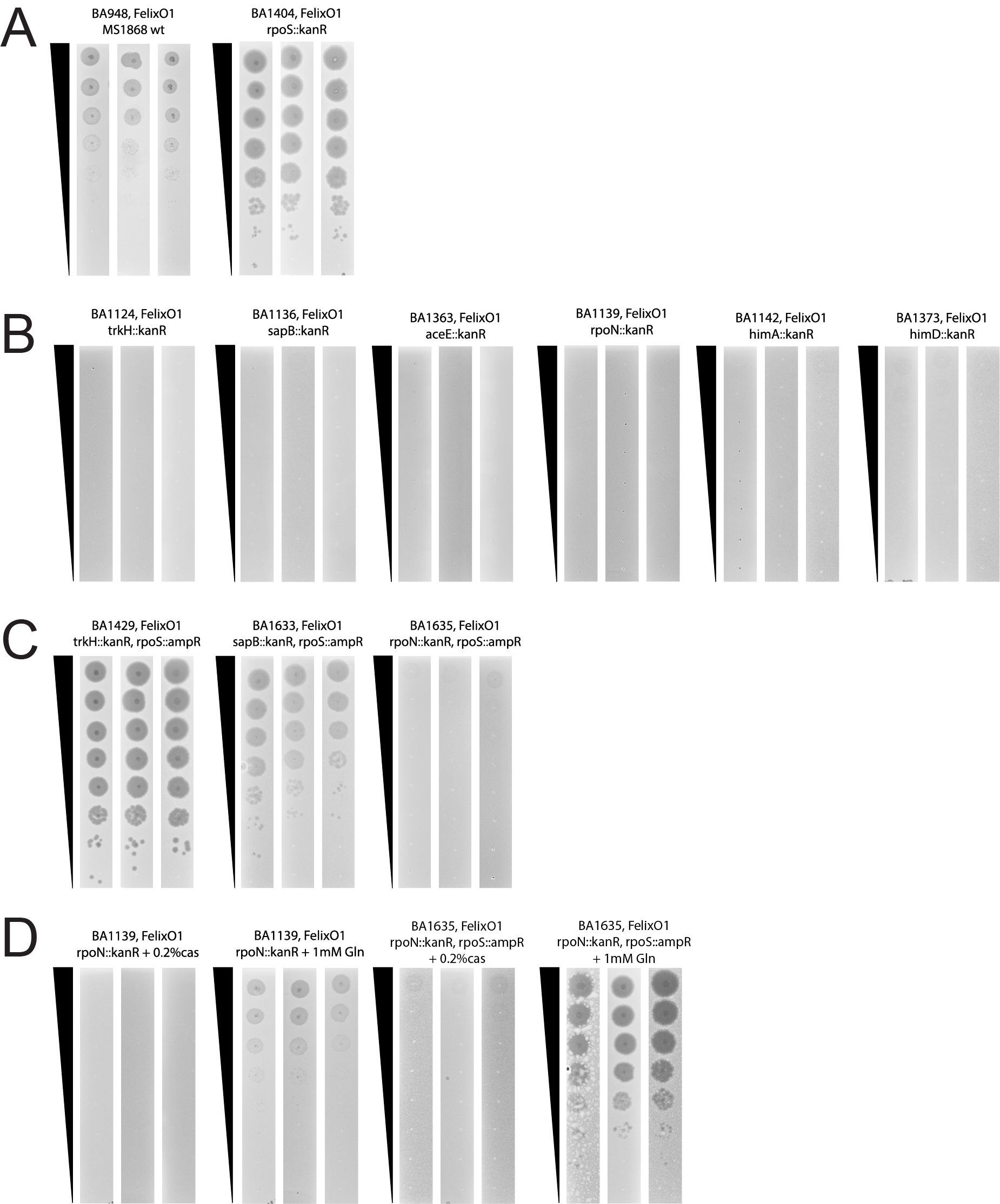

**Figure S20. FelixO1 Followups for Cross-Resistance Mutants**

Plaque assays for phage FelixO1 against mutants identified in phage cross-resistance. In all panels, genotypes for the strains are described. Three biological replicates from unique days are shown.

A. FelixO1 against wildtype and *Salmonella* *rpoS* mutant.

B. FelixO1 plated against single deletions of cross-resistant genes.

C. FelixO1 plated against double deletions of cross-resistant genes and *rpoS*.

D. FelixO1 plated against *rpoN* and *rpoN, rpoS* mutants supplemented with casamino acids or glutamine.

*Table S1. Overview of the S.Typhimurium MS1868 RB-TnSeq Library^[1]^*

| **Barcode-Level Statistics** | |
| --- | --- |
| Total Number of Barcodes | 66,996 |
| Barcodes Associated with a Unique Gene | 55,675 |
| Barcodes Analyzed^[2]^ | 49,655 |
| Mean Barcodes per Gene-Used | 14.8 |
| Median Barcodes per Gene-Used | 12.0 |
| **Gene-Level Statistics** | |
| Genes in MS1868^[3]^ | 4610 |
| Genes Mutated | 3759 |
| Genes Mutated - Primary Chromosome | 3670 |
| Genes Mutated - PSLT Plasmid | 89 |
| Genes Without Mutants | 851 |
| Known Genes Without Unique Mapping^[4]^ | 23 |
| Likely Essential Genes^[5]^ | 434 |
| Likely Nonessential Genes^4^ | 380 |
| MS1868-Specific Genes Without Mutants^4^ | 17 |

*Table S2. Known and Unique hits per phage per screen*

Summary of known and new hits per phage per screen. Total host factors reported here are the total number of genes that meet the thresholds reported in S5 Dataset following manual curation. For details concerning how manual curation was performed, see Materials and Methods.

| ***S.* Typhimurium MS1868 RB-TnSeq screen: Total hits 283** | | | |
| --- | --- | --- | --- |
| Phage | Total host factors | Unique host factors | Previously Reported |
| Aji_GE_EIP16 | 34 | 34 | N/A |
| Br60 | NA | NA | NA |
| Chi | 98 | 90 | 8 |
| FelixO1 | 73 | 61 | 12 |
| Ffm | NA | NA | NA |
| P22 | 20 | 6 | 14 |
| Reaper_GE_8C2 | 16 | 16 | N/A |
| S16 | 7 | 4 | 3 |
| Savina_GE_6H2 | 15 | 15 | N/A |
| Shishito_GE_6F2 | NA | NA | N/A |
| SP6 | 20 | 20 | 0 |
| ***E. coli* K-12 BW25113 RB-TnSeq screen: Total hits 18** | | | |
| Phages | Total host factors | Newly Reported | Previously Reported |
| Br60 | 1 | 1 | 0 |
| Ffm | 10 | 3 | 7 |
| Shishito_GE_6F2 | 7 | 7 | N/A |

*Table S3. Primers used in this study.*

| Primer Number | Primer Name | Purpose | Sequence |
| --- | --- | --- | --- |
| oBA1007 | oBA1007_1100R | 16S verification primers | GGGTTGCGCTCGTTG |
| oBA1008 | oBA1008_337F | 16S verification primers | GACTCCTACGGGAGGCWGCAG |
| oBA757 | oBA757_GFF276_KanR_KO_F | Recombineering template ∆trkH::kanR | GTTAGCGATTGAAGAATAATCCCCACCTCATTTTTAAGAATAAAGGAAGCGGCAGAGATGCTGATCCTTCAACTCAGCAAAAGTTCGATT |
| oBA758 | oBA758_GFF276_KanR_KO_R | Recombineering template ∆trkH::kanR | CAATTTCGCGCGTTTGTCCGTCCCGGGTAGAGAAAAGAATCAATGTTTTCACGTGTTACTCCATTAGAGTCCCGTCAAGTCAGCGTAATG |
| oBA769 | oBA769_GFF2689_KanR_KO_F | Recombineering template ∆sapB::kanR | AATGCGTCTTTTGCCGGCGTCTCCCGCGAAAAACACGAAGAGGTGAAAAAACCATGATTACTGATCCTTCAACTCAGCAAAAGTTCGATT |
| oBA770 | oBA770_GFF2689_KanR_KO_R | Recombineering template ∆sapB::kanR | ACGCCAGGCAGTGCGCAGGGTGCCTGGCGGGCGCTTTTCGCTGTATACGCTATCGTAAGGCATACCGAGTCCCGTCAAGTCAGCGTAATG |
| oBA771 | oBA771_GFF2946_KanR_KO_F | Recombineering template ∆rpoN::kanR | CAGACTCTGATAGGGTAGAGGGCTATGCTGCTCTAGCGGGAGAAAACGACTCTGAATATGCTGATCCTTCAACTCAGCAAAAGTTCGATT |
| oBA772 | oBA772_GFF2946_KanR_KO_R | Recombineering template ∆rpoN::kanR | TCAGTAATTTCGACATTATGTCCGGTGATATTGAGCTGCATAGTGTCTTCCTTATCGGTTGGGTCAGAGTCCCGTCAAGTCAGCGTAATG |
| oBA775 | oBA775_GFF3528_KanR_KO_F | Recombineering template ∆himA::kanR | CCAAATGTGTAGAGGCATTAAAAGAGCGATTCCAGGCATCATTGAGGGATTGAACCTATGCTGATCCTTCAACTCAGCAAAAGTTCGATT |
| oBA776 | oBA776_GFF3528_KanR_KO_R | Recombineering template ∆himA::kanR | TACTTTCGGGATGGCAGCGTATCTGCCGCAATACACCCTGATGGATGTTATGCCTGGATCTGATTAGAGTCCCGTCAAGTCAGCGTAATG |
| oBA964 | oBA964_GFF122_aceE_Kan_KO_R | Recombineering template ∆aceE::kanR | TCAACTTCATCTGTCCCGATGTCCGGTACTTTGATTTCGATAGCCATTATTCTTTTACCTCTTAGAGTCCCGTCAAGTCAGCGTAATGCT |
| oBA965 | oBA965_GFF122_aceE_Kan_KO_F | Recombineering template ∆aceE::kanR | GGGACAGGTTCCAGATAACTCAACGTATTAGATAGATAAGGAATACCCCCATGCTGATCCTTCAACTCAGCAAAAGTTCGATTTATTCAA |
| oBA968 | oBA968_GFF2736_tonB_Kan_KO_R | Recombineering template ∆tonB::kanR | AAGTATACCCGCTTACGCCGCCAGCAGGTGATGGTATATTCCTGGCTGGCGGCGCCAGAGATTAGAGTCCCGTCAAGTCAGCGTAATGCT |
| oBA969 | oBA969_GFF2736_tonB_Kan_KO_F | Recombineering template ∆tonB::kanR | TGCATTTAAAATTCAGCTCTGGTTTTTCAACTGAAACGATTATGACTTCAATGCTGATCCTTCAACTCAGCAAAAGTTCGATTTATTCAA |
| oBA970 | oBA970_GFF4489_fepA_Kan_KO_R | Recombineering template ∆fepA::kanR | AATGAGATGTCAGCATCGTTTTTGCCAATTCCCTCCCCGAATGAGGGAGGGAAGGTTGCCATCAGAGTCCCGTCAAGTCAGCGTAATGCT |
| oBA971 | oBA971_GFF4489_fepA_Kan_KO_F | Recombineering template ∆fepA::kanR | TTGACGGGCGCTTTGGCTTATGTGGCTAAAGAAAAGCAGGATATACAATGAACCTGATCCTTCAACTCAGCAAAAGTTCGATTTATTCAA |
| oBA972 | oBA972_GFF4489_rpoS_Kan_KO_R | Recombineering template ∆rpoS::kanR | CAGAAGACAAACGGTAAAAAAAAGGCCAGTCGACAGACTGGCCTTTTTTTGACAAGGGTACTTAGAGTCCCGTCAAGTCAGCGTAATGCT |
| oBA973 | oBA973_GFF4489_rpoS_Kan_KO_F | Recombineering template ∆rpoS::kanR | AGGCTTTGACTTGCTAGTTCCGTCAAGGGATCACGGGTAGGAGCCACCTTTTGCTGATCCTTCAACTCAGCAAAAGTTCGATTTATTCAA |
| oBA978 | oBA978_GFF3669_ihfB_Kan_KO_R | Recombineering template ∆himD::kanR | CGTATAAATGAAAAAAGCACCCTGACGGTGCTTTTTTCGGGTTCAAGTTTTGCGTTAAAACTTAGAGTCCCGTCAAGTCAGCGTAATGCT |
| oBA979 | oBA979_GFF3669_ihfB_Kan_KO_F | Recombineering template ∆himD::kanR | ATCAATCTCACGGCTGCAGCCAATTTGCCTTTAAGGAACCGGAGGAATCATGCTGATCCTTCAACTCAGCAAAAGTTCGATTTATTCAAC |
| oBA1030 | oBA1030_GFF4489_rpoS_ampR_KO_F | Recombineering template ∆rpoS::ampR | ACTTGCTAGTTCCGTCAAGGGATCACGGGTAGGAGCCACCTTTTGATAACCCTGATAAATGCTTCAATAATATTGAAAAAGGAAGAGTAT |
| oBA1031 | oBA1031_GFF4489_rpoS_ampR_KO_R | Recombineering template ∆rpoS::ampR | AGACAAACGGTAAAAAAAAGGCCAGTCGACAGACTGGCCTTTTTTTGACAAGGGTACTTATTACCAATGCTTAATCAGTGAGGCACCTAT |
| oBA1073 | oBA1073_STM2232_oafA_KanR_KO_F | Recombineering template ∆oafA::kanR | ATCCATTATCTTAATTTCGTCTTGTGTGGCACCTTGGAATTATAGGTAAAAAATGCTGATCCTTCAACTCAGCAAAAGTTCGATTTATTC |
| oBA1074 | oBA1074_STM2232_oafA_KanR_KO_R | Recombineering template ∆oafA::kanR | ATAAAATAATTTGCATTATTGTTGTAGTTTTATAAAATAAAAAGAGGGGCAAGCCCCTCTGTTTAGAGTCCCGTCAAGTCAGCGTAATGC |
| oBA779 | oBA779_GFF276_Nterm_check_F | Verification primers: trkH locus | GCCATTAACCGAATATACTTTGCAGTGTGAG |
| oBA1192 | oBA1192_trkH_DS_R | Verification primers: trkH locus | GCGCCAATCACTACGCGATCATAGC |
| oBA785 | oBA785_2689_Nterm_check_F | Verification primers: sapB locus | GGAGAAAGAGCTGCCGATACTGCC |
| oBA1193 | oBA1193_sapB_DS_R | Verification primers: sapB locus | CAGACCGGCGCATCCATACAGAC |
| oBA786 | oBA786_2946_Nterm_check_F | Verification primers: rpoN locus | GAACGCGCCTATATCGTGAGCCA |
| oBA1194 | oBA1194_rpoN_DS_R | Verification primers: rpoN locus | CTGTTCAAGTTTGGCGAATTTTGTCGTCAC |
| oBA788 | oBA788_3528_Nterm_check_F | Verification primers: ihfA locus | GCGGAGGGTTATAAGAGCCTCGC |
| oBA1195 | oBA1195_ihfA_DS_R | Verification primers: ihfA locus | CGGGCAAAAGTCAGCATGTTATCCATTC |
| oBA985 | oBA985_GFF122_aceE_UP_F | Verification primers: aceE locus | GTTTATCGAAGAGATTATGCTGGACAGAAGCC |
| oBA986 | oBA986_GFF122_aceE_DN_R | Verification primers: aceE locus | GGACTTCCATAGAGGCTTTGTCGCC |
| oBA989 | oBA989_GFF2736_tonB_UP_F | Verification primers: tonB locus | GCGAGTTGGCATTGTCCTGAGCG |
| oBA990 | oBA990_GFF2736_tonB_DN_R | Verification primers: tonB locus | CGATCCGGACGGTAAACCTCGC |
| oBA991 | oBA991_GFF4489_fepA_UP_F | Verification primers: fepA locus | CCACCAAAAAGTGACCCGATAATTTCCGTC |
| oBA992 | oBA992_GFF4489_fepA_DN_R | Verification primers: fepA locus | GCAGGTCGTGTTCGCGAAAGCT |
| oBA993 | oBA993_GFF4788_rpoS_UP_F | Verification primers: rpoS locus | GGGCAAAAAATCGCTACTATGGGTAGCAC |
| oBA994 | oBA994_GFF4788_rpoS_DN_R | Verification primers: rpoS locus | CGCGAACACTATCCACAAGCGTTTC |
| oBA999 | oBA999_GFF3669_ihfB_UP_F | Verification primers: ihfB locus | GCAATGGCTGAAGCATTCAAAGCAGC |
| oBA1000 | oBA1000_GFF3669_ihfB_DN_R | Verification primers: ihfB locus | GACCGTCGTTATCTTCATAGACACCTCCC |
| oBA1095 | oBA1095_STM2232_oafA_check_F | Verification primers: oafA locus | GTCGTCGCGTGGCGAAACG |
| oBA1096 | oBA1096_STM2232_oafA_check_R | Verification primers: oafA locus | GGCGACGAACTGGCGCAGTAC |

*Table S4. Plasmids used in this study.*

| Plasmid | Selection | Source |
| --- | --- | --- |
| pSIM5 | CmR | Gift from Donald Court |

*Table S5. Strains used in this study.*

| Strain Number | Genotype | Source | Notes |
| --- | --- | --- | --- |
| BA948 | *Salmonella enterica* serovar Typhimurium MS1868 | Richard Calendar | *S.* Typhimurium LT2 *(leuA414(Am) Fels2- hsdSB(r-m+)* *(Graña, Youderian, and Susskind 1985)* |
| BA1171 | *Escherichia coli* BW25113 | *E. coli* Genetic Stock Center |  |
| BA1150 | *S.* Typhimurium MS1868 + pSIM5 | This Study |  |
| BA1124 | *S.* Typhimurium MS1868, *∆trkH::kanR* | This Study |  |
| BA1136 | *S.* Typhimurium MS1868, *∆sapB::kanR* | This Study |  |
| BA1139 | *S.* Typhimurium MS1868, *∆rpoN::kanR* | This Study |  |
| BA1142 | *S.* Typhimurium MS1868, *∆himA::kanR* | This Study |  |
| BA1366 | *S.* Typhimurium MS1868, *∆tonB::kanR* | This Study |  |
| BA1368 | *S.* Typhimurium MS1868, *∆fepA::kanR* | This Study |  |
| BA1372 | *S.* Typhimurium MS1868, *∆himD::kanR* | This Study |  |
| BA1404 | *S.* Typhimurium MS1868, *∆rpoS::kanR* | This Study |  |
| BA1460 | *S.* Typhimurium MS1868, *∆oafA::kanR* | This Study |  |
| BA1417 | *S.* Typhimurium MS1868, *∆rpoS::ampR* | This Study |  |
| BA1429 | *S.* Typhimurium MS1868, *∆trkH::kanR, ∆rpoS::ampR* | This Study |  |
| BA1633 | *S.* Typhimurium MS1868, *∆sapB::kanR, ∆rpoS::ampR* | This Study |  |
| BA1635 | *S.* Typhimurium MS1868, *∆rpoN::kanR, ∆rpoS::ampR* | This Study |  |
| BA1232 | SGSC1412 - *S.* Typhimurium LT2 wt | Kenneth Sanderson, *Salmonella* Genetic Stock Center |  |
| BA1234 | SGSC68 - SL428 | Kenneth Sanderson, *Salmonella* Genetic Stock Center | *rfc-458*, semi-rough (Ra with 1 side chain unit) |
| BA1236 | SGSC163 - SL1306 | Kenneth Sanderson, *Salmonella* Genetic Stock Center | *galE503*, Rc |
| BA1238 | SGSC225 - TSL3770 (a) | Kenneth Sanderson, *Salmonella* Genetic Stock Center | *rfa*+, smooth |
| BA1240 | SGSC227 - SL3748 (a) | Kenneth Sanderson, *Salmonella* Genetic Stock Center | *rfaI432*, Rb3 |
| BA1242 | SGSC228 - SL3749 (a) | Kenneth Sanderson, *Salmonella* Genetic Stock Center | *rfaL446*, Ra |
| BA1244 | SGSC229 - SL3750 (a) | Kenneth Sanderson, *Salmonella* Genetic Stock Center | *rfaJ417*, Rb2 |
| BA1246 | SGSC230 - SL3789 (a) | Kenneth Sanderson, *Salmonella* Genetic Stock Center | *rfaF511*, Rd2 |
| BA1248 | SGSC231 - SL3769 (a) | Kenneth Sanderson, *Salmonella* Genetic Stock Center | *rfaG471*, Rd1 |
| BA1250 | SGSC258 - SL1102 | Kenneth Sanderson, *Salmonella* Genetic Stock Center | *rfaE543*, Re |
| BA1252 | SGSC356 - SL4807 | Kenneth Sanderson, *Salmonella* Genetic Stock Center | *rfaB707*, mixture of Rc + smooth |
| BA1254 | SGSC358 - SL3600 | Kenneth Sanderson, *Salmonella* Genetic Stock Center | *rfaD657, Re* |
| BA1256 | SGSC389 - SL733 | Kenneth Sanderson, *Salmonella* Genetic Stock Center | *rfaK953*, Rb1 |
| BA1258 | SA1377 (b) | Kenneth Sanderson, *Salmonella* Genetic Stock Center | *rfaC630*, Re |
| BA1260 | SA1627(his-642) | Kenneth Sanderson, *Salmonella* Genetic Stock Center | *rfb-his* deletion, Ra |

**Dataset S1 (separate file).** Gene-level statistics for genes within the MS1868 RB-TnSeq Library.

**Dataset S2 (separate file).** Overview of RB-TnSeq Experiments.

**Dataset S3 (separate file).** Raw RB-TnSeq Data.

**Dataset S4 (separate file).** High Fitness RB-TnSeq Data.

**Dataset S5 (separate file).** RNA-Seq StringTie Data.

**Dataset S6 (separate file).** RNA-Seq DEseq2 Data.

**Dataset S7 (separate file).** Genome Sequences for Newly Isolated Phages.

**Dataset S8 (separate file).** Gene-level variation observed amongst SARA isolates.

**Dataset S9 (separate file).** Prophage-level variation observed amongst SARA isolates.
